# Metabolic profile discriminates and predicts *Arabidopsis* susceptibility to virus under field conditions

**DOI:** 10.1101/2020.11.21.392688

**Authors:** Bernadette Rubio, Olivier Fernandez, Patrick Cosson, Thierry Berton, Mélodie Caballero, Fabrice Roux, Joy Bergelson, Yves Gibon, Valérie Schurdi-Levraud

## Abstract

As obligatory parasites, plant viruses alter host cellular metabolism. There is a lack of information on the variability of virus-induced metabolic responses among genetically diverse plants in a natural context with daily changing conditions. To decipher the metabolic landscape of plant-virus interactions in a natural setting, one hundred and thirty-two and twenty-six accessions of *Arabidopsis thaliana* were inoculated with *Turnip mosaic virus* (TuMV), in two field experiments over 2 years. The accessions were phenotyped for viral accumulation, above-ground biomass, targeted and untargeted metabolic profiles. The accessions revealed quantitative response to the virus, from susceptibility to resistance. Susceptible accessions accumulate primary and secondary metabolites upon infection, at the cost of hindered growth. Orthogonal Partial Least Squares-Discriminant Analysis (OPLS-DA) revealed that the primary metabolites sucrose, glucose and glutamate discriminate susceptible and resistant accessions. Twenty-one metabolic signatures were found to significantly accumulate in resistant accessions whereas they maintained their growth at the same level as mock-inoculated plants without biomass penalty.

Metabolic content was demonstrated to discriminate and to be highly predictive of the susceptibility of inoculated *Arabidopsis*. The PLS coefficient estimated in the training data set reveals, after cross-validation, a correlation of 0.61 between predicted and true viral accumulation. This study is the first to describe the metabolic landscape of plant-virus interactions in a natural setting and its predictive link to susceptibility. It reveals that, in this undomesticated species and in ecologically realistic conditions, growth and resistance are in a permanent conversation and provides new insights on plant-virus interactions.

## Introduction

Plant health is of primary importance to improve and secure food supply for an increasing human population. Plant viruses represent the major taxonomic group of emergent pathogens of plants (Anderson *et al*., 2004), and viral infection is one of the most alarming biotic threats due to the impact of climate change on the spatial and temporal distribution of vectors and viruses (Bebber *et al*., 2013; Jones, 2016). Compared to other plant pathogens, viruses are particularly unpredictable and difficult to combat. In this context, an understanding of the response of plants to viral infection has great importance in sustainable agricultural solutions.

The genus *Potyvirus*, to which the *Turnip mosaic virus* (TuMV) belongs, is one of the largest genera among plant viruses, causing considerable economic damage in vegetable and fruit crops worldwide (Adams *et al*., 2011). The completion of the viral multiplication and movement cycle results from a complex interplay between virus- and host-encoded factors that can have profound impacts on plant fitness. To invade plants, those obligatory parasites have developed tactics to reroute host cellular functions and components for their own benefits. Whatever the outcome of the interaction – compatibility, leading to disease, or incompatibility, leading to resistance – massive reprogramming of metabolism is observed. Indeed, as viruses perturb and exploit the host’s carbon (Stare *et al*., 2015) and nitrogen (Fernandez-Calvino *et al*., 2014) metabolism to make their own compounds, demand for energy within the plant increases to sustain viral multiplication, systemic spread and defense responses (Biemelt & Sonnewald, 2006).

To date, viral impacts on plant metabolism have been mainly studied in experiments conducted in controlled conditions and on a limited diversity of host genotypes (typically a few genotypes). In these conditions, respiration (Shalitin *et al*., 2002), photosynthetic efficiency (Kangasjärvi *et al*., 2012) and carbon partitioning (Fernandez-Calvino *et al*., 2014) were shown to be modified. For example, in a susceptible interaction between tobacco and *Potato virus Y* (PVY), both an increase of soluble carbohydrates and a decrease of photosynthesis were observed 4 days after inoculation (dai) (Handford and Carr, 2007). These observations were confirmed by the demonstration that potato leaves exhibit a decrease in sugar levels one day after PVY infection, with a subsequent increase in both inoculated and systemic leaves a few days later (Stare *et al*., 2015; Kogovsek *et al*., 2016). Similarly, inoculation of susceptible hosts with *Cucumber mosaic virus* (CMV) results in a localized reduction in starch accumulation as a consequence of altered carbohydrate metabolism at viral infection sites (Tecsi *et al*., 1996; Shalitin *et al*., 2002), although starch accumulates to high levels in systemically infected leaves (Handford & Carr, 2007). Increases in free amino acid content have also been demonstrated in terms of total or individual amino acids, as well as in polyamines, in a variety of systems (reviewed in Llave, 2016; Pilar Lopez-Gresa *et al*., 2012; Fernandez-Calvino *et al*., 2014). Viruses interfere with fatty acids, structural components of intracellular membranes in which replication can take place (Heaton & Randall, 2011). They also exploit transport systems in order to invade cells in systemic tissues away from the initial site of infection. This is achieved through an interaction between viral proteins and components of the long distance transport machinery like phloem proteins (Wang, 2015). Like primary metabolism, secondary metabolism is strongly impacted due to its participation in multiple defense signaling cascades (Piasecka *et al*., 2015). Numerous compounds, including hormones, are involved in defense (Verma *et al*., 2016; Palukaitis *et al*., 2017). Among them, glucosinolates and phytoalexins play significant roles in defense against a range of pathogens (Kliebenstein, 2004).

In a compatible interaction, the outcome of viral colonization can include symptoms such as stunting, chlorosis or necrosis depending on the pathosystem. Incompatible interactions trigger plant resistance and defense signaling that involve the action of antimicrobial components and specific defense proteins (Piasecka *et al*., 2015). These defense responses are now widely acknowledged to involve a trade-off in model plants, with the cost of resistance generating a negative impact on plant fitness (Tian et al, 2003; Denance *et al*., 2013; Neilson *et al*., 2013; Huot *et al*., 2014; Lozano-Duran & Zipfel, 2015; Karasov *et al*., 2017; Albrecht & Argueso, 2017). Similarly, in crop plants, high levels of resistance are often associated with yield penalties (Bergelson & Purrington 1996; Brown, 2002).

While extremely informative, these studies performed under controlled optimal conditions reduce environmental effects and increase the likelihood of finding strong relationships between metabolite levels variations (Fernandez *et al*., 2016). Thereby, it obscures the complexity and the variability of metabolic responses among genetically diverse plants in a natural context where they have to face multiple stresses. Even in main crop species, large-scale metabolic profiling of large field-grown populations of genetically diverse accessions remains rare (Schauer et al., 2006; Obata et al., 2015; Melandri et al., 2019).

In *Arabidopsis thaliana*, metabolite profiling of large populations of natural accessions or inbred lines has allowed the identification of descriptor sets of metabolites that are predictive of biomass (Meyer et al., 2007; Sulpice et al., 2010; Steinfath et al., 2010) and physiological traits such as freezing tolerance (Korn et al., 2010) and herbivore resistance (Züst et al., 2012). But, none of these studies addressed the growth and metabolic response to biotic stress in a changing environment.

Here, following the modern standards of ecological genomics (Bergelson & Roux, 2010), we aimed at improving our understanding of the virus-induced reprogramming of metabolism through deciphering variation in the metabolic landscape of the natural *Arabidopsis thaliana*/*Turnip mosaic virus* (TuMV) pathosystem in growth conditions close to environmental reality. To do so, we set up two experiments in the field and explored targeted and untargeted metabolic profiles on 132 and 26 accessions, in 2014 and 2015 respectively. These accessions spanned quantitative responses ranging from high susceptibility to full resistance. This work aimed to (i) decipher the trade-offs among responses to TuMV, metabolism and growth in field conditions in a large set of *Arabidopsis* accessions, (ii) characterize the metabolic disturbances for a large set of accessions, and (iii) identify discriminant metabolic biomarkers of susceptibility/resistance to viruses in *A. thaliana*.

## Material and Method

### Plant Material

A worldwide collection of natural accessions of *Arabidopsis thaliana* were used in our experiments (Horton *et al*., 2012). One hundred thirty-two accessions were challenged with TuMV during the first field experiment in 2014 as described in Rubio *et al*. (2019) and then a subset of 26 accessions were selected for a second field experiment in 2015. This subset was selected based on their OD value following TuMV inoculation (see description of “Quantification of viral accumulation”) (listed in Table S1) to represent extreme phenotypes from highly susceptible to resistant. To prevent misidentifying accessions as resistant due to inefficient mechanical inoculation, only accessions presenting a resistance phenotype in all common garden experiments were selected as resistant accessions (Rubio et al. 2019).

### Virus Material

*Turnip mosaic virus* isolate UK1 (Jenner & Walsh, 1996) was routinely propagated by mechanical inoculations on turnip plants *Brassica rapa* L. ssp *rapa* NA FR 490001 provided by the BraCySol germplasm center (Ploudaniel, France). To prepare the inoculum, three-week old turnip plants were mechanically inoculated. Symptoms appeared two weeks later. Young symptomatic leaves of five-week-old turnip were then collected to produce the inoculum.

### Experimental Design and Growth conditions

Two common garden experiments were conducted in 2014 (N=132 *A. thaliana* accessions) and 2015 (N=26 accessions). Each experiment was organized in a randomized complete block design (RCBD). In 2014, the experiment contained three blocks with one replicate per accession per block whereas the experiment performed in 2015 contained four blocks with four replicates per accession per block. As described in Rubio *et al*. (2019), seedling trays of 40 wells were used. Seeds were sown on 24 March 2014 and 23 March 2015 in professional horticultural soil (106Scope, PletrAcom, Arles, France) under a cold-frame glasshouse without additional light or heating to ensure homogeneity of germination. At three weeks of age, plants were acclimatized under an opened tunnel before their transfer to the common garden. Plants were inoculated during the acclimation step just before their transfer. Soil of the common garden had been tilled so that seedling trays could be slightly buried. Because the bottoms of the wells were pierced, roots were able to reach the soil. Climate data were recorded over the duration of each experiment (Figure S1, A and B). The analysis of climatic data across the three growing environments (cold frame greenhouse, then tunnel, then field) revealed no significant difference between years 2014 and 2015 (Figure S1, C, D and E) for temperatures, rainfall and PAR (photosynthetically active radiation).

### TuMV inoculation procedure and harvest

One gram of fresh turnip leaves was ground in three volumes of disodium phosphate (Na_2_HPO_4_.12H2O) 30mM and 0.2 % diethyldithiocarbamic acid (DIECA). The inoculum was clarified through 10 min centrifugation at 13,000 g. Supernatant was recovered and maintained at 4°C until Arabidopsis’ inoculation, which was performed when plants were 4 weeks old and at a 8-10 leaf stage, corresponding to 1.09, 1.10 Boyes stage (Boyes *et al*., 2001). Four young expanded leaves of each plant were mechanically inoculated with 20µL of inoculum with carborandum added on each leaf. Ten minutes after inoculation, plants were rinsed with water. Mock treatments, in which plants were treated exactly as inoculated plants except for the absence of the virus, were also included. The viral concentration of the TuMV inoculum was quantified after inoculation by quantitative PCR (Rubio *et al*., 2019). There were no significant differences between the viral concentrations of the inocula used in the 2014 (0.073 ng/µL) and 2015 (0.103 ng/µL) common garden experiments (Wilcoxon test, p-value = 0.0636).

According to previous experiments by Rubio *et al*. (2019), samples were collected 13 days after inoculation (dpi) at the very beginning of the onset of the first symptoms. The harvest was done in the morning, at the same time for each year of experiment. All rosette leaves above the inoculated leaves (systemic leaves) were collected in vials from Zinsser Analytic® (Eschborn, Germany). Samples were deep-frozen, ground and stored at -80°C.

### Quantification of viral accumulation

Viral accumulation was estimated for each individual plant using 100 mg of *A. thaliana* powder in a semi-quantitative double antibody sandwich assay (DAS-ELISA) with a commercial anti-potyvirus monoclonal antibody kit (Agdia-Biofords, Evry, France). The reaction of the substrate (p-nitrophenyl phosphate) was followed at 405 nm. Optical densities (OD) were calculated by removing the mean OD value from the healthy *A. thaliana* Col-0 control and normalized using a Col-0 positive control deposited on each ELISA plate. As determined from the ODs of the *A. thaliana* Col-0 positive controls, and in order to avoid overflow, the 15 min measurements were retained. For each accession, we classified its degree of resistance or susceptibility based on the average of the ODs obtained across replicates (Table S1). Resistant accessions do not accumulate virus and present an OD equal or lower than the mean of the OD of the healthy *A. thaliana* Col-0 control (OD=0.088 SD 0.01063). Susceptible accessions have an OD value that exceeds the healthy *A. thaliana* Col-0 control. The category to which each accession belongs is listed in Table S1, along with its OD values. Mock-inoculated plants were confirmed to be potyvirus-free.

### Sample processing

Three biological replicates were prepared in 2014 and four biological replicates were prepared in 2015. In 2015, a biological replicate was composed by pooling up to four plants after a successful infection as measured by DAS ELISA. Aliquots of about 17 mg of fresh powder were weighted in 1.1-mL Micronic tubes and used in the targeted metabolite analysis. Samples were then lyophilized and aboveground dry mass determined with an analysis balance. Aliquots of about 10 mg were weighed in 1.1-mL Micronic tubes and used in the untargeted metabolite analysis.

### Targeted Metabolite analysis

Metabolites were extracted in a final volume of 650µL, twice with 80 % (v/v) ethanol – HEPES/KOH 10mM (pH6) and once with 50 % (v/v) ethanol – HEPES/KOH 10mM (ph6) (Geigenberger *et al*., 1996). Chlorophyll content was determined immediately after the extraction (Arnon, 1949). Glucose, fructose, sucrose (Stitt & Quick, 1989) malate, fumarate (Nunes-Nesi *et al*., 2007), glutamate (Zhang *et al*., 2015) and amino acids (Bantan-Polak *et al*., 2001) were determined in the supernatant. Starch (Hendriks *et al*., 2003) and protein (Bradford, 1976) contents were determined on the pellet resuspended in 100 mM NaOH. Analyses were performed in 96-well microplates using Starlet pipetting robots (Hamilton), and absorbance was read in MP96 microplate readers (SAFAS). **Data treatments**. Metabolite data were normalized by dividing each measure by the value of a biological standard sample corresponding to a non-inoculated *A. thaliana* Col-0 sample harvested at the same time as the other samples. The final concentration of each metabolite is the average of the replicates of each accession. **Statistical analysis**. All principal component analysis (PCA), parametric and nonparametric statistical tests were performed using R version 3.4.0. Statistical significance was set at P < 0.05. Partial Least Square regression (PLS) and Orthogonal Partial Least-Squares regression and discriminant analysis (OPLS-DA) were performed using the R packages mixOmics (Rohart *et al*., 2017) and PLS (Wehrens & Mevik, 2007).

### Untargeted Metabolic analysis

Untargeted metabolite measurements were conducted on the subset of 26 accessions in the common garden experiment in 2015. Extraction was realized using a Starlet pipetting robot (Hamilton) with an extraction buffer composed of 80 % (v/v) ethanol and 0.1 % (v/v) formic acid, using methyl vanillate as an internal standard (50µg/mL). All samples extracted were filtrated through a Multiscreen Solvinert 96-well filter plate (Merck Millipore). **Quality control (QC)**. 25 µL of each sample were mixed together to generate a pooled quality control sample (QC). QC samples were analyzed every 10 injections to monitor and correct changes in the instrument response. Solvent blank samples (80% ethanol in water – 0.1 % formic acid) were also analyzed in-between the other samples. **Liquid chromatography**. Liquid chromatography was performed on a Dionex UHPLC Ultimate 3000 (Thermo Scientific, MA, USA). Chromatographic separation was carried out in reverse-phase mode on a Gemini C18 column (2 x 150 mm; 3µm, Phenomenex, CA, USA) equipped with a Gemini C18 guard column (2 x 4 mm, Phenomenex, CA, USA). The mobile phase was composed of milliQ water with 0.1% formic acid (solvent A) and 100% Acetonitrile (solvent B) for a total run time of 18 min. The flow rate was 0.3 mL.min^-1^ and the column was heated to 30°C. The autosampler temperature was maintained at 4°C and the injection volume was 5 µL. **Mass spectrometry**. The UHPLC system was coupled with a LTQ-Orbitrap Elite mass spectrometer (Thermo Scientific, MA, USA). A heated electrospray interface was used and analyses were performed in positive mode. Acquisition was performed in full scan mode with a resolution of 240 000 FWHM in the scan range of m/z 50-1000. Data were recorded using Xcalibur software (Thermo Scientific, MA, USA) and extracted with XCMS. **Data processing and statistical analysis**. Data were converted to mzXML file format. Peak picking and alignment were performed using XCMS in R (Smith *et al*., 2006). XCMS-parameters were optimized as described by Patti and collaborators (Patti *et al*., 2012). XCMS-data processing results in a data matrix which contains peak intensities that are a unique combination of retention-time and median m/z ratio. To exclude system-peaks (impurities in the measurement-system, visible in blanks) as well as poorly detected metabolic features, filter steps were performed based on QCs and blanks as described in TextS1. Statistical analysis was done in R version 3.4.0 and also using the web application BioStatFlow version 2.7.7. Thus, OPLS-DA were performed using the 505 features of the untargeted metabolic analysis combined with the 10 primary metabolites assayed by targeted methods.

## Results

### Susceptible accessions accumulate primary metabolites upon infection, at the cost of hindered growth, whereas resistant accessions grow with limited changes

The experiment conducted in 2014 showed that twenty genotypes out of one hundred and thirty inoculated were considered fully resistant as no virus was detected (Rubio et al., 2019; Table S1). The remaining accessions fell within a range of susceptibility, from the least susceptible (OD > 0.088 healthy *A. thaliana* Col-0 control OD; Table S1) to the most susceptible (OD=0.896; Table S1).

Aboveground dry biomass was measured on mock-inoculated and TuMV-inoculated plants. Aboveground dry biomass was significantly lower for the 110 inoculated susceptible accessions compared to those susceptible accessions when treated only with a mock solution (Wilcoxon-Mann Whitney test, *p*.*value*=0.039; Figure 1A), indicating that infection led to a decrease in the growth of susceptible plants. There was no significant difference between the aboveground dry biomass of the 20 resistant inoculated accessions compared to their mock-inoculated counterparts (Wilcoxon-Mann Whitney test, *p*.*value*=0.1274, Figure 1A). This result was confirmed for the 18 susceptible and 8 resistant accessions included among the 132 (Table S1) accessions used in the 2015 field experiment (Figure 1B; Wilcoxon-Mann Whitney test, *p*.*value*=3.44 E^-8^ of susceptible accessions; *p*.*value*=0.1040 for resistant accessions). Spearman correlations confirmed that viral accumulation was negatively correlated with dry aboveground biomass in both years (Table S2 A and B).

**Figure 1.**
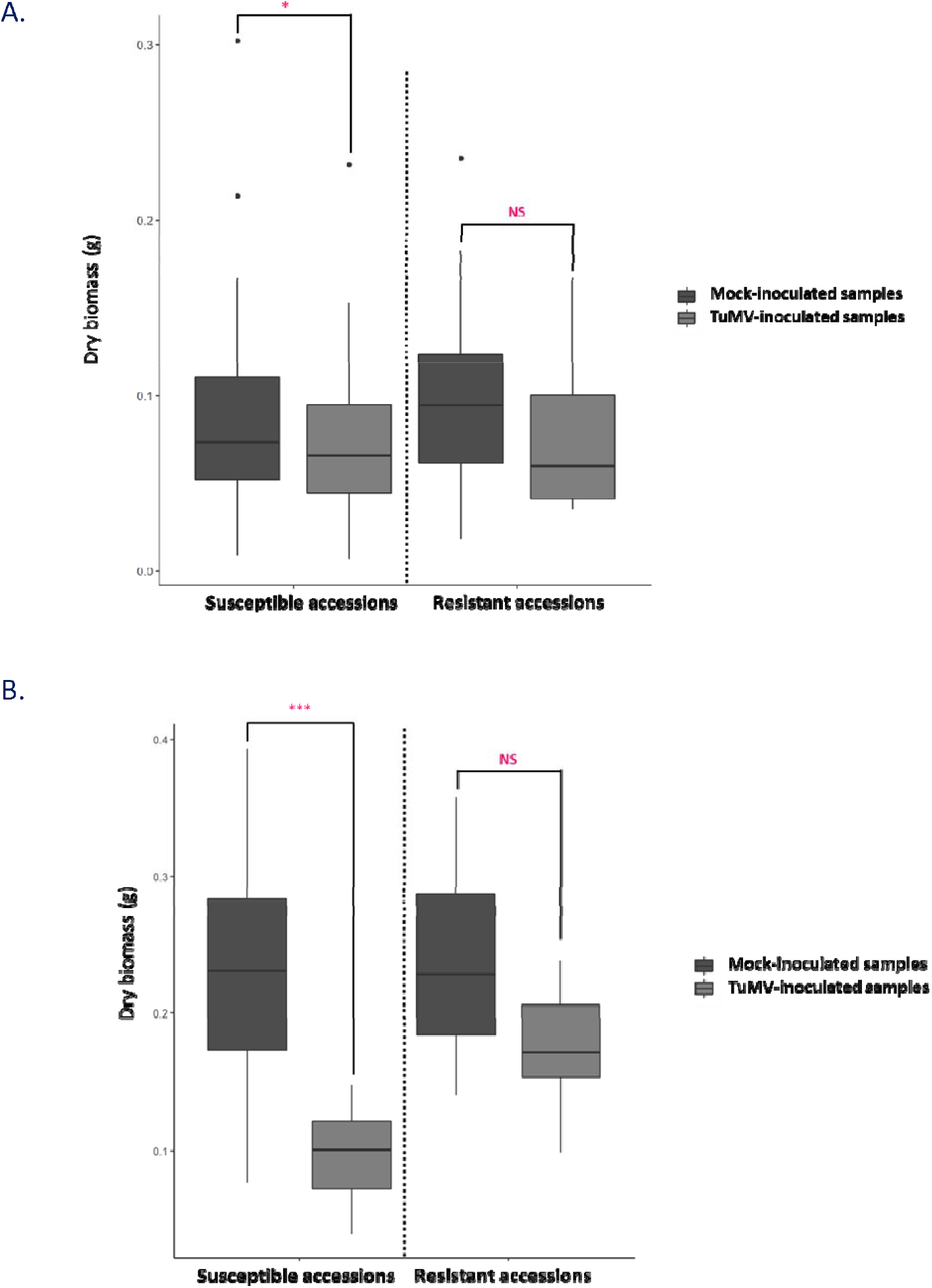
Dry biomass for mock-inoculated and TuMV-inoculated susceptible and resistant accessions measured at 13 days after TuMV inoculation on (A) 130 *A. thaliana* and on (B) 26 accessions in the ‘2014’ and ‘2015’ field experiment, respectively. Statistical comparisons (Wilcoxon-Mann Whitney test) on dry biomass were performed between TuMV and mock-inoculated samples on both categories of accessions. Signif.codes: ‘*’ *P* < 0.05; ‘***’ *P* < 0.001

Ten key metabolic traits were measured for each accession in each of the years. In mock-inoculated samples, a significant positive correlation was found between malate, fumarate, starch, glucose, fructose, sucrose and aboveground dry biomass whereas a negative significant correlation was found between amino acids, proteins, glutamate and aboveground dry biomass (Table S2 B).

In neither year was the chlorophyll content in TuMV-inoculated susceptible accessions significantly reduced, confirming that sampling occurred prior to macroscopic symptoms of chlorosis (Figure 2A and 2B). In 2014, the 110 susceptible inoculated accessions accumulated significant larger amounts of glutamate and fructose compared to both resistant inoculated accessions and mock-inoculated accessions (Figure 2A). Similarly, in 2015, the 18 TuMV-inoculated susceptible accessions exhibited a significant accumulation of 8 out of 10 primary metabolites (i.e. amino acids, glutamate, malate, fumarate, starch, glucose, fructose and sucrose), when compared to either the 8 resistant inoculated accessions or the set of mock inoculated controls (Figure 2B). These results were supported by significant positive correlations between viral accumulation expressed as OD and eight out of ten primary metabolites content - amino acids, proteins, glutamate, fumarate, starch, glucose, fructose and sucrose (Table S2 A and B) but significantly negative correlations between the metabolites content and the above ground biomass (Table S2B). In addition to maintaining their growth, the inoculated resistant accessions exhibited similar amounts of primary metabolites relative to those observed for mock-inoculated controls.

**Fig. 2.**
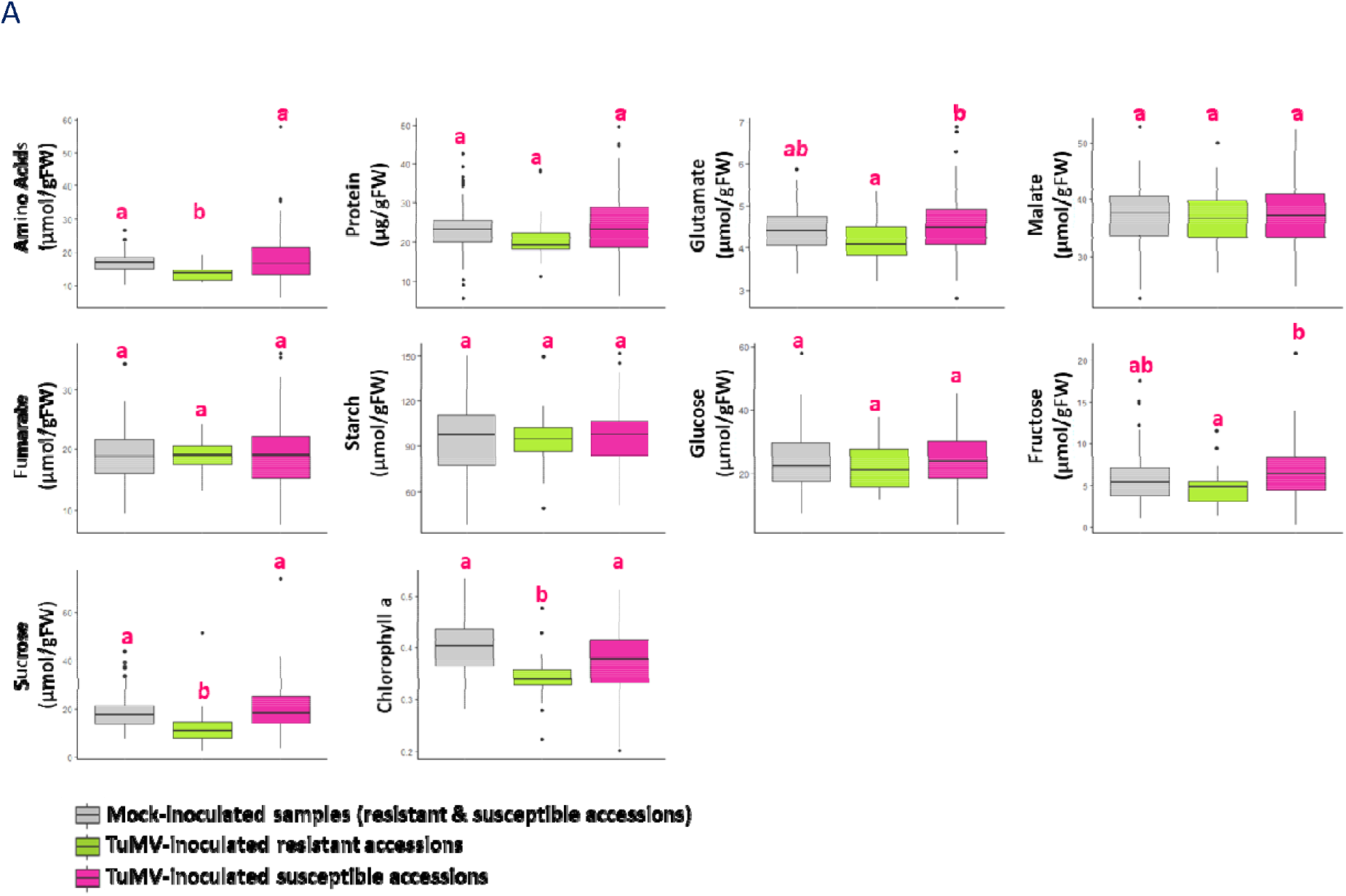
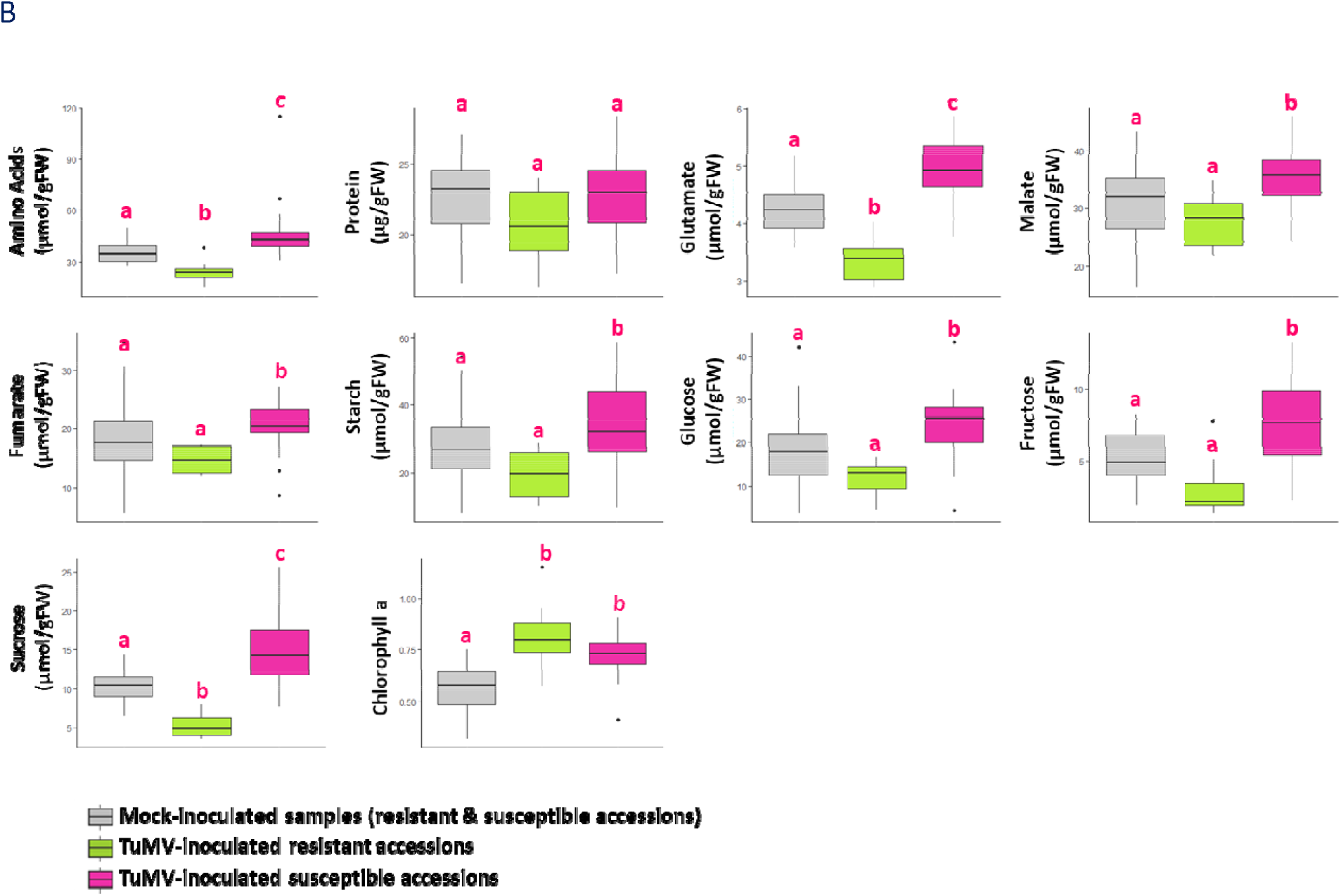
Comparisons of primary metabolic content in mock-inoculated and TuMV-inoculated resistant and susceptible *A. thaliana* accessions in the two-year field experiment. **A. *A. thaliana* accessions of ‘2014’ field experiment** **B. *A. thaliana* accessions of ‘2015’ field experiment** Mock-inoculated samples are represented by grey boxplot. TuMV inoculated resistant accessions are represented by green boxplot and susceptible accessions by pink boxplots. Statistical analyzes were performed on each of the 10 primary metabolites (Wilcoxon-Mann Whitney or Student test according to the normality of the data). For each metabolite, plots with the same letter are not significantly different at *P* = 0.05.

### Metabolic content discriminates inoculated A. thaliana susceptible and resistant accessions

Targeted metabolism analysis performed on 132 genotypes in 2014 and 26 genotypes in 2015 does not distinguish between susceptible and resistant genotypes when mock-inoculated (Figure 3 IA and IIA). The broad, non-targeted analysis conducted by UHPLC-LTQ Orbitrap on the set of 26 genotypes in 2015 also fails to distinguish between susceptible and resistant genotypes when mock-inoculated (Figure 4A).

**Figure 3.**
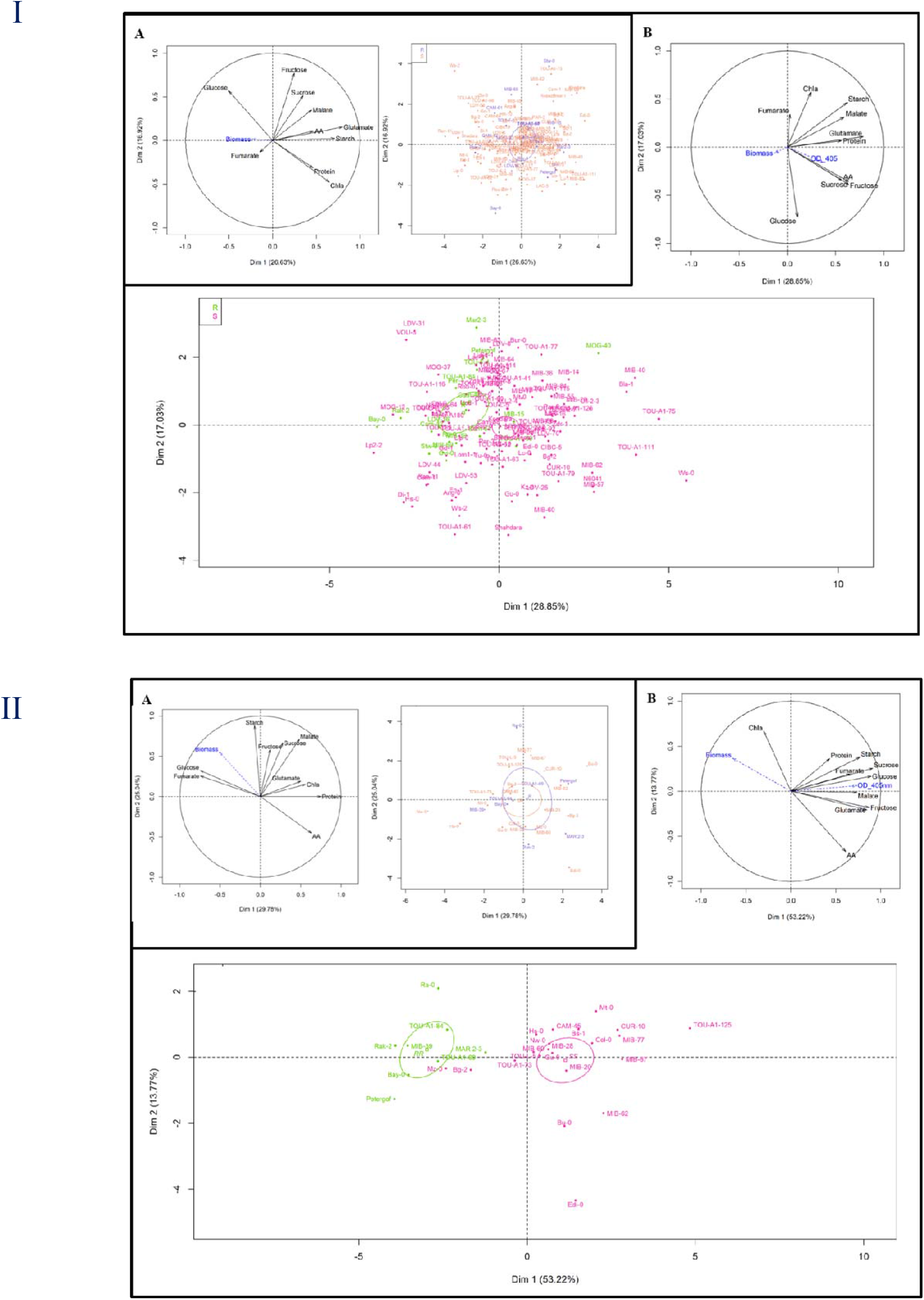
Principal component analysis (PCA) performed on (I) 130 *A. thaliana* accessions inoculated by TuMV and mock-inoculated in the ‘2014’ field experiment and on (II) 26 *A. thaliana* accessions inoculated by TuMV and mock-inoculated in the ‘2015’ field experiment. **A**. PCA performed with 10 primary metabolic traits measured at 13 days on mock-inoculated *A. thaliana* accessions. The two major components that together accounted for 43.55% of the variance in 2014 and 50.55% in 2015 have been plotted. Dry biomass (Biomass) and viral accumulation (OD_405nm) were considered as explanatory traits (in blue on the variable factor map). Score plot of resistant (in blue) and susceptible (in orange) accessions. The confidence ellipses around the centroid of individuals are represented. **B**. PCA performed with 10 primary metabolic traits measured at 13 days after TuMV inoculation The two major components that accounted for 45.88% of the variance in 2014 and 68.14% in 2015 have been plotted. Dry biomass (Biomass) and viral accumulation (OD_405nm) were considered as explanatory traits (in blue on the variable factor map). Score plot of resistant (in green) and susceptible (in pink) accessions. The confidence ellipses around the centroid of individuals are represented. AA = Amino Acids, Chla = Chlorophyll a.

**Figure 4.**
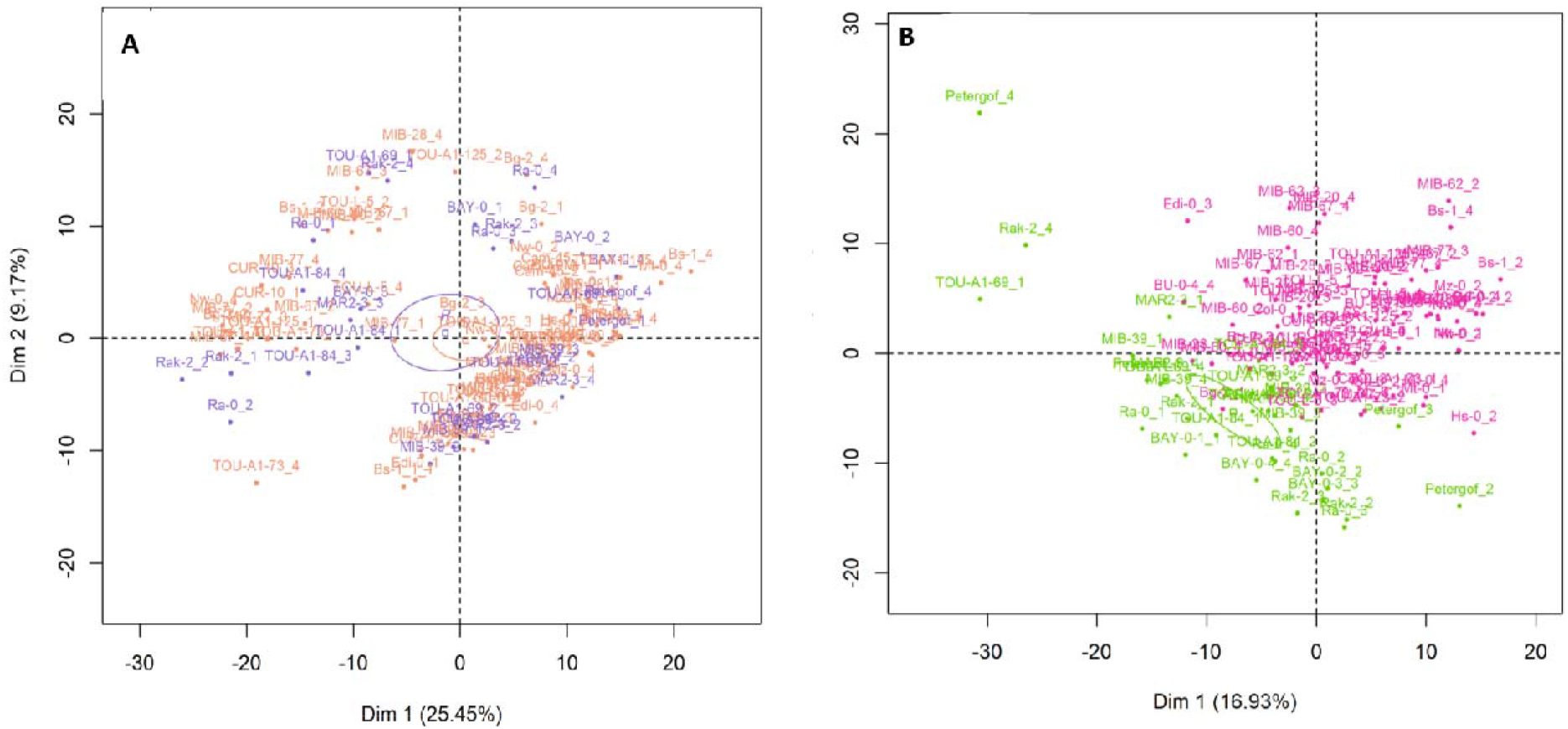
Principal component analysis performed on the 505 metabolic signatures (m/z) measured on 26 *A. thaliana* accessions in the ‘2015’ field experiment. **A**. PCA performed on the 505 metabolic signatures (m/z) measured by UHPLC-LTQ Orbitrap on 26 *A. thaliana* mock-inoculated accessions. The two major components that together accounted for 34.62% of the variance. Resistant and susceptible accessions are in blue and orange, respectively. **B**. PCA performed on the 505 metabolic signatures (m/z) measured by UHPLC-LTQ Orbitrap on 26 *A. thaliana* TUMV-inoculated accessions. The two major components accounted for 26.05% of the variance. Resistant and susceptible accessions are in green and pink, respectively.

The composition of metabolites differed between TuMV-inoculated plants (Figure 3 IB and IIB) and mock-inoculated plants (Figure 3 IA and IIA). Given that the vast majority of metabolic change occurred in susceptible plants, this response leads to a clear differentiation of metabolic content between susceptible and resistant accessions that have been inoculated with TuMV in both years (Figure 3 IB and IIB; Figure S2). The first two axes of the PCA explained 45.88% and 63.14% of the overall variance of the metabolic profiles in inoculated samples in 2014 and 2015 respectively. In 2015, the first axis of the PCA, in particular, discriminates between susceptible and resistant accessions (54.31% of the explained variance) even though these accessions are indistinguishable in terms of metabolic composition when mock-inoculated (Figure 3 IA and IIA).

To analyze the pattern of infection and its relationship to a larger set of metabolites, including secondary metabolites, we conducted an untargeted metabolic analysis by UHPLC-LTQ Orbitrap on the set of 26 accessions showing a contrasting response to TuMV in the 2015 field experiment. This analysis captured a total of 505 workable metabolic signatures (m/z), corresponding primarily to non-polar secondary metabolites. As previously observed with targeted primary metabolism, there was no difference between susceptible and resistant accessions when mock-inoculated (Figure 4A). In contrast, when inoculated with TuMV, principal component analysis on the metabolic variables clearly separated two groups of accessions according to their susceptibility to the TuMV (Figure 4B).

### The identification of metabolic predictors reveals 21 metabolic signatures significantly accumulated in resistant accessions

An Orthogonal Partial Least Squares-Discriminant Analysis (OPLS-DA) was then performed to maximize the variation between the two groups of accessions and determine the most significant variables contributing to this variation, i.e. VIPs (Variable Importance in the Projection). OPLS-DA analysis was carried out using the primary and secondary metabolite data. The quality of the model was validated by the Q^2^ parameter (goodness-of-prediction parameter) with a value of 0.846, thereby showing high predictive capabilities (Figure S3).

The OPLS-DA analysis performed between the mock- and TuMV-inoculated samples revealed 63 common discriminant metabolic variables (Table S3), most of which (58/63) accumulated in TuMV-inoculated samples and particularly in susceptible accessions (54/58; Table S3). It is worth noting that four VIPs (433;700, 361;491, 512;491 and 64;395) are significantly found in resistant accessions.

To examine which metabolic variables strongly contribute to the OPLS-DA model in TuMV-inoculated susceptible and resistant accessions, variables were ranked according to their VIP values (Table 1). This ranking confirmed the major role of some primary metabolites, including sucrose, glucose, fructose, glutamate, amino acids and fumarate, in the response of susceptible accessions. Of the 140 VIP values identified by OPLS-DA, 119 accumulated to higher levels in susceptible accessions and 21, among which the 4 VIPs (VIP 433;700, VIP 361;491, VIP 512;491 and VIP 64;395) are found, accumulated to higher levels in resistant accessions (Table 1). The fold change susceptible/resistant reached up to 45 times whereas the fold change resistant/susceptible reached up to 244 times. It is very interesting to highlight two VIPs, VIP 324;184 and VIP 433;482 that are significantly accumulated in resistant accessions when inoculated with a fold-change of 106.8 and 244, respectively, compared to the susceptible accessions.

**Table 1.** List of the Variable Importance in the Projection (VIPs) identify by OPLS-DA analysis performed on TuMV-inoculated resistant and susceptible twenty-six accessions in 2015. For each VIP, comparisons between resistant (R) and susceptible (S) accessions were done. The fold change was calculated for each VIP. Primary metabolites are light-grey highlighted. Metabolites that accumulate significantly more in resistant accessions are at the bottom of the table.

| VIP OPLS-DA<sup>1</sup> | VIP values | m/z<sup>2</sup> | rt<sup>3</sup> | Resistant vs Susceptible<br>metabolic contents | Fold<br>change |
| --- | --- | --- | --- | --- | --- |
| Sucrose | 2.2126449 | NA | NA | S > R ***<sup>4</sup> | 2.89 |
| 303;463 | 1.9733637 | 303.133385 | 463.27 | S > R *** | 3.69 |
| 356;452 | 1.9579956 | 356.120178 | 452.352 | S > R *** | 2.17 |
| 219;311 | 1.9395195 | 219.101071 | 311.134 | S > R *** | 4.81 |
| 533;392 | 1.9257616 | 533.154889 | 391.765 | S > R *** | 4.98 |
| 332;430 | 1.8980522 | 332.131528 | 430.481 | S > R *** | 3.78 |
| 116;100 | 1.8668978 | 116.070147 | 100.163 | S > R *** | 6.05 |
| 205;242 | 1.8374938 | 205.096695 | 242.121 | S > R *** | 2.51 |
| 103;242 | 1.7986954 | 103.041077 | 242.089 | S > R *** | 2.86 |
| 302;506 | 1.7961978 | 302.101728 | 506.314 | S > R *** | 5.76 |
| 255;544 | 1.7691068 | 255.112248 | 543.799 | S > R *** | 9.96 |
| 221;216 | 1.7482809 | 221.091514 | 215.744 | S > R *** | 3.13 |
| 385;210 | 1.7379538 | 385.105549 | 210.083 | S > R *** | 45.77 |
| Glucose | 1.7341573 | NA | NA | S > R *** | 2.49 |
| 175;402 | 1.7128412 | 175.147752 | 401.831 | S > R *** | 3.77 |
| 903;319 | 1.7101545 | 903.276945 | 318.889 | S > R *** | 8.48 |
| 503;391 | 1.6946581 | 503.190232 | 391.457 | S > R *** | 2.51 |
| 343;345 | 1.6930327 | 343.116792 | 345.319 | S > R *** | 2.08 |
| Fructose | 1.6832118 | NA | NA | S > R *** | 2.49 |
| 474;107 | 1.674786 | 474.217815 | 107.298 | S > R *** | 2.68 |
| 315;448 | 1.6735275 | 315.133358 | 447.821 | S > R *** | 2.39 |
| 209;488 | 1.669006 | 209.153094 | 488.35 | S > R *** | 2.7 |
| 209;414 | 1.6625752 | 209.153104 | 413.756 | S > R *** | 2.14 |
| 212;284 | 1.6568285 | 211.559242 | 283.891 | S > R *** | 1.56 |
| 543;99 | 1.6447422 | 543.132047 | 98.5221 | S > R *** | 5.84 |
| 124;346 | 1.6383986 | 124.075207 | 346.354 | S > R *** | 1.92 |
| 370;302 | 1.6097363 | 370.148552 | 301.519 | S > R *** | 28.49 |
| 203;204 | 1.5909149 | 203.084422 | 203.918 | S > R *** | 8.24 |
| Glutamate | 1.5902017 | NA | NA | S > R *** | 1.47 |
| 430;333 | 1.5812953 | 430.169872 | 332.878 | S > R *** | 3.3 |
| 331;329 | 1.5779633 | 331.117148 | 328.817 | S > R *** | 1.58 |
| 226;406 | 1.5540098 | 226.106582 | 406.369 | S > R *** | 7.97 |
| 151;359 | 1.548784 | 151.075013 | 359.377 | S > R *** | 1.78 |
| 449;319 | 1.545091 | 449.106284 | 318.949 | S > R *** | 1.75 |
| 757;319 | 1.5374453 | 757.217144 | 319.13 | S > R *** | 1.69 |
| 270;587 | 1.5342534 | 270.133029 | 587.091 | S > R *** | 43.68 |
| 315;370 | 1.5236406 | 315.133322 | 370.476 | S > R *** | 3.93 |
| 162;402 | 1.5107457 | 162.054596 | 402.281 | S > R *** | 1.9 |
| 221;230 | 1.5031416 | 221.120712 | 230.244 | S > R *** | 1.74 |
| 394;517 | 1.4896806 | 394.204469 | 517.072 | S > R *** | 17.28 |
| 302;407 | 1.4864035 | 302.101313 | 406.697 | S > R *** | 7.16 |
| 482;101 | 1.4749114 | 482.107244 | 101.274 | S > R *** | 1.79 |
| Amino Acids | 1.4618098 | NA | NA | S > R *** | 1.88 |
| 355;306 | 1.4506109 | 355.101534 | 306.288 | S > R *** | 3.29 |
| 191;416 | 1.4478404 | 191.142656 | 416.467 | S > R *** | 2.49 |
| 193;324 | 1.4445315 | 193.125245 | 323.817 | S > R *** | 1.46 |
| 149;360 | 1.4367383 | 149.095721 | 360.18 | S > R *** | 1.66 |
| 642;445 | 1.43456 | 642.253866 | 445.078 | S > R *** | 2 |
| 321;760 | 1.4189617 | 321.114081 | 760.442 | S > R *** | 3.7 |
| 182;466 | 1.4146411 | 182.080769 | 465.796 | S > R *** | 4.21 |
| 305;210 | 1.408059 | 305.086054 | 210.166 | S > R *** | 2.24 |
| 348;319 | 1.3963139 | 348.2736 | 318.945 | S > R *** | 3.61 |
| 164;496 | 1.3753555 | 164.070296 | 496.307 | S > R *** | 2.22 |
| 165;424 | 1.3719233 | 165.127004 | 424.14 | S > R *** | 1.54 |
| 317;464 | 1.3699949 | 317.101381 | 463.65 | S > R *** | 1.25 |
| 191;363 | 1.3599054 | 191.069895 | 362.643 | S > R *** | 2.76 |
| 219;486 | 1.3401209 | 219.101009 | 486.477 | S > R *** | 2.7 |
| 105;700 | 1.3304992 | 105.06927 | 700.287 | S > R *** | 1.59 |
| 367;358 | 1.3232566 | 367.10133 | 358.491 | S > R *** | 1.69 |
| 201;344 | 1.3189902 | 201.054173 | 343.832 | S > R *** | 1.71 |
| 189;325 | 1.3123658 | 189.127006 | 324.754 | S > R *** | 1.74 |
| 146;242 | 1.3107417 | 146.059816 | 241.843 | S > R *** | 2.31 |
| 133;296 | 1.3066147 | 133.06438 | 295.68 | S > R *** | 2.65 |
| 179;604 | 1.3057044 | 179.106264 | 603.961 | S > R *** | 1.64 |
| 374;322 | 1.3047581 | 374.143633 | 321.703 | S > R *** | 1.77 |
| 373;453 | 1.2900149 | 373.127302 | 452.6 | S > R *** | 1.81 |
| 302;382 | 1.2818991 | 302.041251 | 381.707 | S > R *** | 1.54 |
| 161;442 | 1.2743927 | 161.095747 | 441.797 | S > R *** | 1.79 |
| 164;375 | 1.2688136 | 164.070227 | 375.012 | S > R *** | 2.95 |
| 109;359 | 1.2670868 | 109.064191 | 359.34 | S > R *** | 1.92 |
| 386;278 | 1.2559975 | 386.219789 | 277.549 | S > R *** | 3.48 |
| 391;774 | 1.2553695 | 391.244577 | 774.477 | S > R *** | 2.29 |
| 409;491 | 1.2512226 | 409.168874 | 490.545 | S > R *** | 27.12 |
| 291;259 | 1.2414233 | 291.180846 | 259.063 | S > R *** | 4.78 |
| 192;462 | 1.2266011 | 192.041337 | 462.37 | S > R *** | 1.27 |
| 107;370 | 1.218738 | 107.084913 | 370.001 | S > R *** | 1.39 |
| 420;276 | 1.2167175 | 419.694896 | 275.797 | S > R *** | 1.77 |
| 193;373 | 1.1889052 | 193.085573 | 373.266 | S > R *** | 2.04 |
| 379;402 | 1.1794454 | 379.094921 | 402.311 | S > R *** | 2.88 |
| 181;464 | 1.1755653 | 181.085582 | 463.916 | S > R *** | 1.42 |
| 181;328 | 1.1704302 | 181.085601 | 327.894 | S > R *** | 1.68 |
| 80;395 | 1.1675747 | 80.0488023 | 395.289 | S > R *** | 1.57 |
| 373;344 | 1.1603378 | 373.127271 | 343.531 | S > R *** | 1.54 |
| 627;299 | 1.1556143 | 627.15471 | 299.369 | S > R *** | 2.8 |
| 96;389 | 1.1547774 | 96.0801348 | 389.004 | S > R *** | 1.74 |
| 195;453 | 1.1541351 | 195.064751 | 452.614 | S > R *** | 1.55 |
| 79;396 | 1.1518191 | 79.0409732 | 395.754 | S > R *** | 1.52 |
| 105;327 | 1.1496278 | 105.069244 | 326.898 | S > R *** | 1.72 |
| 210;465 | 1.1465056 | 210.111814 | 464.799 | S > R *** | 2.44 |
| 335;506 | 1.1433297 | 335.126504 | 506.441 | S > R *** | 3.71 |
| 86;126 | 1.1144812 | 86.0957928 | 126.246 | S > R *** | 2.09 |
| 396;257 | 1.1063852 | 396.185419 | 256.51 | S > R *** | 1.43 |
| 611;354 | 1.1020274 | 611.158212 | 353.603 | S > R *** | 1.34 |
| 178;191 | 1.1016652 | 178.089265 | 191.013 | S > R *** | 3.43 |
| 169;496 | 1.1007391 | 169.049224 | 495.5 | S > R *** | 3.67 |
| 162;249 | 1.0978873 | 162.054721 | 249.101 | S > R *** | 1.5 |
| 396;348 | 1.0917369 | 396.114946 | 347.739 | S > R *** | 4.14 |
| 103;389 | 1.0860577 | 103.053575 | 389.444 | S > R *** | 2 |
| 521;325 | 1.0838985 | 521.201084 | 325.328 | S > R *** | 2.52 |
| 133;126 | 1.0715649 | 133.104827 | 126.347 | S > R ** | 2.32 |
| 209;426 | 1.0709285 | 209.15308 | 425.939 | S > R *** | 1.98 |
| 162;357 | 1.0685924 | 162.054596 | 356.879 | S > R *** | 1.74 |
| 201;451 | 1.0634552 | 201.054251 | 451.123 | S > R *** | 1.75 |
| 527;460 | 1.0557011 | 527.103495 | 460.095 | S > R *** | 2.02 |
| Fumarate | 1.0539467 | NA | NA | S > R *** | 1.38 |
| 393;416 | 1.0524847 | 393.187545 | 415.816 | S > R *** | 1.71 |
| 212;774 | 1.0524681 | 212.094368 | 774.047 | S > R *** | 2.76 |
| 402;420 | 1.0483873 | 402.161979 | 419.597 | S > R ** | 1.87 |
| 404;388 | 1.038078 | 404.227082 | 388.155 | S > R ** | 1.31 |
| 464;367 | 1.0348068 | 464.248021 | 366.989 | S > R *** | 1.97 |
| 225;371 | 1.0338844 | 225.147985 | 370.798 | S > R *** | 1.57 |
| 103;284 | 1.0330701 | 103.05364 | 283.823 | S > R *** | 3.26 |
| 227;789 | 1.0226386 | 227.163338 | 788.506 | S > R *** | 1.61 |
| 222;216 | 1.0155496 | 221.601711 | 215.822 | S > R ** | 1.33 |
| 367;342 | 1.0107207 | 367.153077 | 341.908 | S > R *** | 1.96 |
| 302;463 | 1.0063676 | 302.101889 | 463.078 | S > R *** | 1.49 |
| 309;325 | 1.0028313 | 309.115614 | 324.659 | S > R ** | 1.44 |
| 244;325 | 1.0006993 | 244.096223 | 324.677 | S > R ** | 1.41 |
| 351;184 | 2.0467196 | 351.006221 | 184.222 | R > S *** | 2.88 |
| 432;184 | 1.8309047 | 431.970682 | 183.776 | R > S *** | 3.44 |
| 137;131 | 1.6182679 | 136.930742 | 131.31 | R > S *** | 1.39 |
| 512;491 | 1.5629621 | 512.127487 | 491.367 | R > S *** | 2.54 |
| 394;108 | 1.547668 | 394.200084 | 108.419 | R > S *** | 1.93 |
| 337;573 | 1.5096853 | 337.292489 | 572.917 | R > S *** | 1.64 |
| 299;181 | 1.3961487 | 299.098193 | 181.363 | R > S *** | 22.3 |
| 181;139 | 1.2934886 | 181.052538 | 138.934 | R > S *** | 1.94 |
| 324;184 | 1.2633811 | 323.988853 | 183.797 | R > S *** | 106.8 |
| 280;184 | 1.2289246 | 280.084227 | 183.768 | R > S *** | 5.9 |
| 361;491 | 1.2178747 | 361.091906 | 490.564 | R > S *** | 2.07 |
| 155;132 | 1.1408996 | 154.941358 | 131.541 | R > S *** | 1.33 |
| 433;482 | 1.1139622 | 433.112069 | 481.588 | R > S *** | 244 |
| 433;700 | 1.1093528 | 433.240637 | 700.261 | R > S *** | 1.82 |
| 64;395 | 1.0910062 | 63.9336655 | 395.047 | R > S *** | 3.6 |
| 256;668 | 1.0892571 | 256.079504 | 667.874 | R > S *** | 2.12 |
| 460;740 | 1.0755049 | 460.268938 | 740.005 | R > S ** | 1.42 |
| 449;442 | 1.0685439 | 449.1069 | 442.311 | R > S *** | 6.17 |
| 244;130 | 1.0487138 | 243.942484 | 130.167 | R > S *** | 1.35 |
| 347;240 | 1.038776 | 347.159419 | 239.659 | R > S ** | 1.48 |
| 86;184 | 1.0146328 | 86.05939 | 184.278 | R > S *** | 1.87 |

<sup>1</sup> When undetermined, VIP are identified through m/z/rt values. VIP values are classified in decreasing order.
<sup>2</sup> mass to charge ratio
<sup>3</sup> retention time
<sup>4</sup> The significance was assessed through a Wilcoxon test at \*\*\* $P < 0.001$ , \*\* $0.001 < P < 0.01$

A complementary analysis on the dataset obtained with the 26 accessions to test whether viral accumulation can be predicted from metabolic data, Partial Least Square (PLS) analysis was performed on the optical density values corresponding to the viral accumulation measured by DAS ELISA for each accession (Figure 5). The PLS coefficient estimated in the training data set reveals, after cross-validation, a correlation of 0.61 between predicted and true viral accumulation, confirming the high predictive power of metabolic composition for TuMV susceptibility. Among the best 50 VIP-PLS values, 44 were found common to the VIP values obtained with the OPLS-DA analysis (Table S4). In both analyses, the primary metabolites sucrose, glucose and glutamate were found to discriminate susceptible and resistant accessions, ranking among the best VIP values (Table S4). Six common VIPs (VIP 394;108, VIP 351;184, VIP 324;184, VIP 280;184, VIP 432;184 and VIP 86;184) detected by the 2 analyses are found accumulated more significantly in resistant accessions.

**Figure 5.**
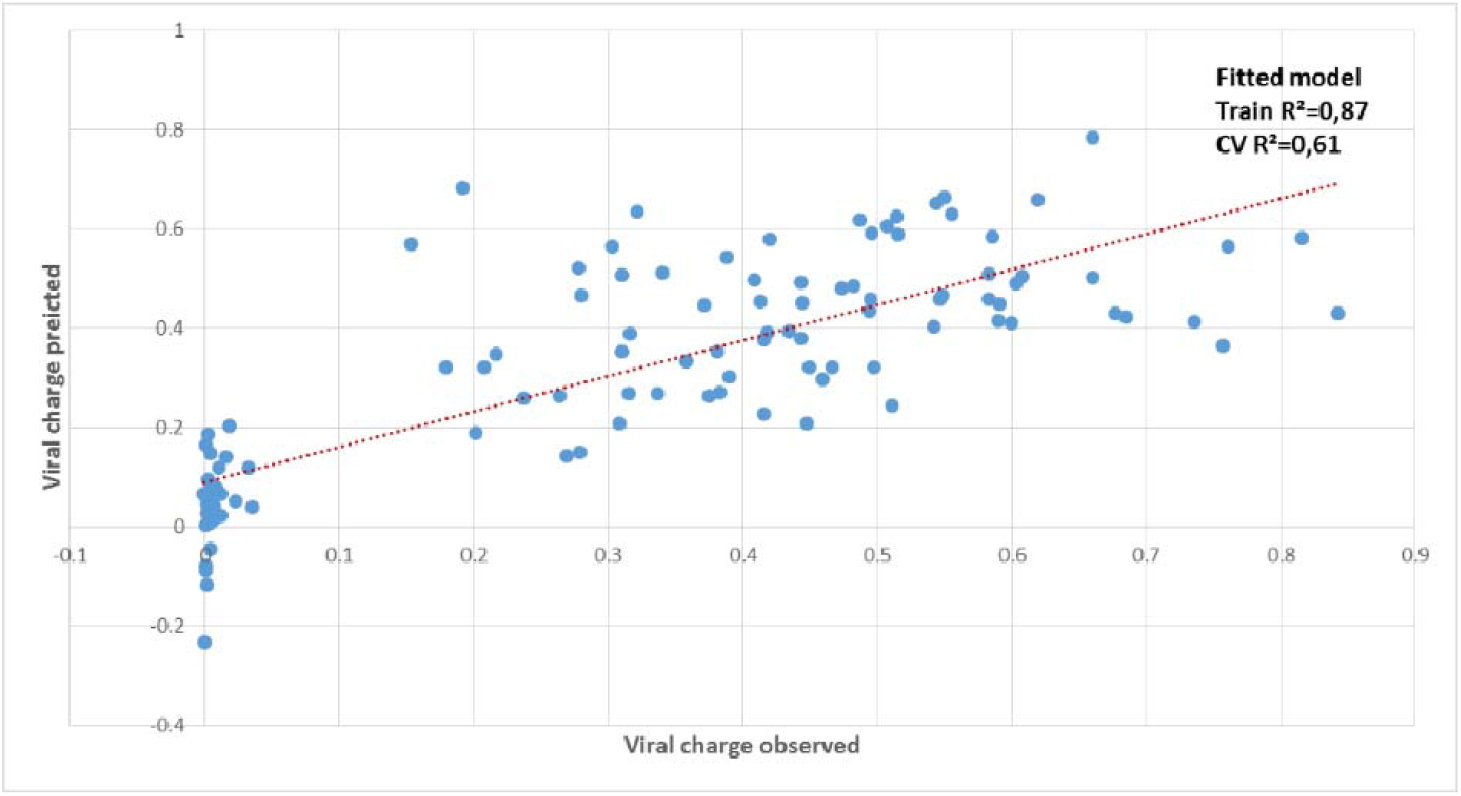
Prediction of viral accumulation by the metabolite matrix on 26 *A. thaliana* accessions in the ‘2015’ field experiment. The metabolite matrix is composed by 10 primary metabolic traits and 505 metabolic signatures (m/z). The replicates of the 26 *A. thaliana* accessions are represented by the blue dots. The dashed linear red line represents the exact prediction.

## Discussion

In this study, we characterized the metabolic response of *A. thaliana* to its natural viral pathogen, *Turnip mosaic virus* (TuMV). This study is unusual in studying the metabolic response of a wide variety of accessions to an important naturally occurring virus on *A. thaliana* (Pagan *et al*., 2010) and its close relative, *A. halleri* (Kamitani *et al*., 2016; 2018), in natural settings. Moreover, in order to mimic ecological reality, this study was conducted for two consecutive years in semi-field conditions, in which plants are subjected to daily and complex environmental changes. It has been shown that the responses of plants to multiple stresses are sometimes very different than those observed under single stress conditions (Rasmussen *et al*., 2013). In field and natural environments experiments, quality and intensity of light, temperature, humidity and many other environmental factors typically vary daily and seasonally, suggesting that plants grown under controlled conditions probably express only a fraction of their responses potential. In natural populations of *A. halleri* in central Japan, Honjo *et al*. (2020) and Nagano *et al*. (2019) have shown through transcriptomic analysis performed over 2 seasons that TuMV infections produces distinct defence mechanisms according to the season. This provides strong arguments for studying plants under realistic field conditions (Jänkänpää *et al*., 2012; Annunziata *et al*., 2017).

In neither year of the study could we distinguish the metabolic profile of TuMV-susceptible and TuMV-resistant accessions grown in the absence of TuMV. This suggest that there are no constitutive metabolic patterns associated with resistance or susceptibility. Nevertheless, these mock-inoculated samples provided an opportunity to describe elements of primary metabolism in *A. thaliana* under common garden field conditions. We found that dry biomass was significantly correlated with fumarate concentrations. Fumarate can accumulate to high levels in *A. thaliana* relative to other plant species, suggesting that it likely constitutes a significant fraction of the fixed carbon in *A. thaliana* rosette leaves (Araújo *et al*., 2011). Indeed, the amount of carbon stored in fumarate is similar to that accumulated in starch (Araújo *et al*., 2011). This is perhaps not surprising because fast-growing plant species such as *A. thaliana* contain significantly higher concentrations of organic acids, such as fumarate, compared to slow-growing plants (Chia *et al*., 2000; Araújo *et al*., 2011), especially under high light intensity conditions such as in our field experiments (Chia *et al*., 2000). We also found that dry biomass was correlated to glucose. Both fumarate and glucose reached high levels in fast growing accessions, 23 mM and 19 mM respectively, suggesting that they might be involved in turgor dynamics and thus growth by cellular expansion (Fricke, 2017). In contrast, we detected a negative correlation between dry biomass and protein. Although speculative, this might follow from the fact that lower protein synthesis contributes to increased efficiency of carbon use, because protein synthesis is a costly process (Sajitz-Hermstein & Nikoloski, 2010; Kafri *et al*., 2015). This negative relationship is in agreement with previous observations among a collection of *A. thaliana* accessions grown under greenhouse/ growth chamber and in controlled conditions (Sulpice *et al*., 2013).

Upon viral infection in the field, susceptible accessions were clearly distinguished from the resistant ones by the accumulation of primary metabolites observed in each of the two years of study. Eight out of ten primary metabolic traits, amino acids, glutamate, malate, fumarate, starch, glucose, fructose and sucrose, were strongly associated with viral accumulation. Previous efforts to describe the massive reprogramming of the plant primary metabolism that takes place in response to pathogens has focused on fungi and bacteria (Rojas *et al*., 2014). Here, we add to the limited literature on plant responses to viral infection, which is restricted to controlled laboratory conditions. For example, in a study of the primary metabolic response of *Arabidopsis* to *Tobacco rattle virus* (TRV), Fernandez-Calvino and collaborators (Fernandez-Calvino *et al*., 2014) showed that the susceptible Col-0 accession significantly accumulated sucrose at 8 days’ post inoculation. Amino acids were also globally accumulated in infected plants compared to mock plants whereas neither starch nor fumarate were accumulated in infected plants. Differences in the virus species, the number of accessions observed and the fact that theirs was a laboratory experiment could explain these different results. Second, in a controlled multi-stress experiment (including TuMV, drought and heat), accumulation of soluble sugars was again observed in Col-0 plants (Prasch and Sonnewald, 2013). The strong metabolic disturbances generated in susceptible plants by TuMV infection suggests that the viral infection stimulated the accumulation of major primary metabolites, probably as a result of an imbalance between photosynthesis and growth, which is known to favor oxidative stress (Foyer and Shigeoka, 2011). Susceptible accessions were also found to accumulate a large number of secondary compounds. This result contrasts with that obtained by Likic *et al*. (2014). In their study conducted under controlled conditions on susceptible Col-0 infected with *Cucumber mosaic virus* (CMVsat), Likic *et al*. (2014) showed that the concentration of some flavonoid compounds such as kaempferol in the upper leaves of all infected plants was significantly lower than that of control plants. In our study, in addition to the fact that such an accumulation was apparently ineffective, the synthesis of secondary compounds is expensive in energy and, even if somewhat controlled (Sirikantaramas *et al*., 2007), accumulation could have toxic effects.

Under the field conditions used in our experiments, inoculated resistant accessions were able to grow at the same rate as controls. As biomass is considered as appropriate proxy for fitness under many circumstances (Younginger *et al*., 2017), it seems to indicate that resistance was achieved with no penalties on measured traits. Maintenance of biomass in relation to moderate symptom development was also observed by Kamitani *et al*. (2016; 2018) in wild populations of *A. halleri* naturally infected by TuMV. Unlike susceptible accessions, resistant accessions accumulate a limited number of metabolites in response to the virus, but in much higher quantities than can be observed in susceptible accessions. This contrasts with a study conducted under controlled conditions on the metabolic response of tomato to *Tomato yellow leaf curl virus* (TYLCV) which found that the defense response of resistant lines is effective through the accumulation of many secondary metabolites (Sade *et al*., 2015).

In a previous study by Rubio *et al*., (2019) the genetic architecture of quantitative response of *A. thaliana* to TuMV in a field environment was reported. The genetic architecture of this interaction reveals at least 10 genomic regions involved in the interaction. One of them overlaps with the gene *AT5G18170* that encodes a subunit of a glutamate dehydrogenase, which integrates carbon and nitrogen metabolism (Terce-Laforgue *et al*., 2013), thereby implicating a link between metabolism and response to virus. The high predictive power of metabolic composition for TuMV susceptibility when infected might enable this composition to serve as a biomarker (Fernandez *et al*., 2016), as has been done for resistance to *Fusarium graminearum* in wheat (Cuperlovic-Culf *et al*., 2016) and for susceptibility to esca disease in grape (Felgueiras *et al*., 2010).

The complex molecular network underlying the balance between growth and immunity has been described (Chandran *et al*., 2014; Huot *et al*., 2014; Lozano-Duran & Zipfel, 2015) mainly in terms of opposition. Recent reviews (Kliebenstein, 2016; Karasov *et al*., 2017) have emphasized, however, that in a natural context, growth and immunity are in a constant conversation. Our study supports this alternative model and highlights new results on *Arabidopsis*/virus interaction. In particular, we find that susceptible accessions experience a large accumulation of primary and secondary metabolites with a reduction of growth whereas resistant accessions appear capable of continued growth with a targeted metabolic response. Some compounds as VIP 324;184 and VIP 433;482, that presented high fold-change in resistant accessions compared to susceptible ones are of particular interest. It would be of great interest to characterize these secondary compounds and their biosynthetic pathways, and then to test their involvement in resistance by using pharmacological and/or genetic approaches. It is worth mentioning that anti-phage secondary metabolites molecules, able to block phage replication, have recently been found in *Streptomyces* (Kronheim *et al*., 2018). This would allow for better understanding the complexity of the underlying mechanisms involved in plants’ responses to viruses in the field and to propose new resilient ideotypes (Sulpice *et al*., 2020).

## Supporting information

Rubio et al. Supplemental information

## Acknowledgements

VS-L thanks the Fulbright Scholarship Program for accommodation and research support in Joy Bergelson’s laboratory. The authors are grateful to Thierry Mauduit and Pascal Briard for their help in the management of field experiments. No conflict of interest to declare.

## References

Adams MJ, King AM, Lefkowitz E, Carstens EB. 2011. Part II: The viruses—family Potyviridae. In: Virus Taxonomy—Ninth Rep. Int. Committee on Taxonomy of Viruses. San Diego, CA, U.S.A.: Elsevier, 1069–1090.

Albrecht T, Argueso CT. 2017. Should I fight or should I grow now? The role of cytokinins in plant growth and immunity and in the growth-defence trade-off. Annals of Botany 119: 725–735.

Anderson PK, Cunningham AA, Patel NG, Morales FJ, Epstein PR, Daszak P. 2004. Emerging infectious diseases of plants: pathogen pollution, climate change and agrotechnology drivers. Trends Ecol Evol 19: 535–544.

Annunziata MG, Apelt F, Carillo P, Krause U, Feil R, Mengin V, Lauxmann MA, Köhl K, Nikoloski Z, Stitt M, et al. 2017. Getting back to nature: a reality check for experiments in controlled environments. Journal of Experimental Botany 68: 4463–4477.

Araújo WL, Nunes-Nesi A, Fernie AR. 2011. Fumarate: Multiple functions of a simple metabolite. Phytochemistry 72: 838–843.

Arnon DI. 1949. Copper enzymes in isolated chloroplasts. Polyphenoloxidase in Beta vulgaris. Plant physiology 24: 1.

Bantan-Polak T, Kassai M, Grant KB. 2001. A Comparison of Fluorescamine and Naphthalene-2,3-dicarboxaldehyde Fluorogenic Reagents for Microplate-Based Detection of Amino Acids. Analytical Biochemistry 297: 128–136.

Bebber DP, Ramotowski MAT, Gurr SJ. 2013. Crop pests and pathogens move polewards in a warming world. Nature climate Change DOI:10.1038/NCLIMATE1990.

Bergelson J, Roux F. 2010. Towards identifying genes underlying ecologically relevant traits in Arabidopsis thaliana. Nat Rev Genet 11: 867–879.

Bergelson J, Purrington CB. 1996. Surveying patterns in the cost of resistance in plants. The American Naturalist, 14, 536–558.

Biemelt S, Sonnewald U. 2006. Plant-microbe interactions to probe regulation of plant carbon metabolism. J Plant Physiol 163: 307–318.

Boyes DC, Zayed AM, Ascenzi R, McCaskill AJ, Hoffman NE, Davis KR, Gorlach J. 2001. Growth stage-based phenotypic analysis of Arabidopsis: a model for high throughput functional genomics in plants. Plant Cell 13: 1499–1510.

Bradford MM. 1976. A rapid and sensitive method for the quantitation of microgram quantities of protein utilizing the principle of protein-dye binding. Analytical biochemistry 72: 248–254.

Brown JKM. 2002. Yield penalties of disease resistance in crops. Current Opinion in Plant Biology 5: 339–344.

Chandran D, Rickert J, Huang Y, Steinwand MA, Marr SK, Wildermuth MC. 2014. Atypical E2F Transcriptional Repressor DEL1 Acts at the Intersection of Plant Growth and Immunity by Controlling the Hormone Salicylic Acid. Cell Host & Microbe 15: 506–513.

Chia DW, Yoder TJ, Reiter W-D, Gibson SI. 2000. Fumaric acid: an overlooked form of fixed carbon in Arabidopsis and other plant species. Planta 211: 743–751.

Cuperlovic-Culf M, Wang L, Forseille L, Boyle K, Merkley N, Burton I, Fobert PR. 2016. Metabolic Biomarker Panels of Response to Fusarium Head Blight Infection in Different Wheat Varieties. PlOS One 11: 1–16.

Denance N, Sanchez-Vallet A, Goffner D, Molina A. 2013. Disease resistance or growth: the role of plant hormones in balancing immune responses and fitness costs. Frontiers in Plant Science 4.

Felgueiras ML, Lima MRM, Dias ACP, Gil AM, Graça G, Rodrigues JEA, Barros A. 2010. NMR metabolomics of esca disease-affected Vitis vinifera cv. Alvarinho leaves. Journal of Experimental Botany 61: 4033–4042.

Fernandez-Calvino L, Osorio S, Hernandez ML, Hamada IB, Del Toro FJ, Donaire L, Yu A, Bustos R, Fernie AR, Martinez-Rivas JM, et al. 2014. Virus-Induced Alterations in Primary Metabolism Modulate Susceptibility to Tobacco rattle virus in Arabidopsis. Plant Physiol 166: 1831–1838.

Fernandez O, Urrutia M, Bernillon S, Giauffret C, Tardieu F, Le Gouis J, Langlade N, Charcosset A, Moing A, Gibon Y. 2016. Fortune telling: metabolic markers of plant performance. Metabolomics 12: 158.

Foyer CH, Shigeoka S. 2011. Understanding Oxidative Stress and Antioxidant Functions to Enhance Photosynthesis. Plant Physiology 155: 93–100.

Fricke W. 2017. Turgor Pressure. In: eLS. American Cancer Society, 1–6.

Geigenberger P, Lerchi J, Stitt M, Sonnewald U. 1996. Phloem-specific expression of pyrophosphatase inhibits long distance transport of carbohydrates and amino acids in tobacco plants. Plant, Cell & Environment 19: 43–55.

Handford MG, Carr JP. 2007. A defect in carbohydrate metabolism ameliorates symptom severity in virus-infected Arabidopsis thaliana. J Gen Virol 88: 337–341.

Heaton NS, Randall G. 2011. Multifaceted roles for lipids in viral infection. Trends in Microbiology 19: 368–375.

Hendriks JH, Kolbe A, Gibon Y, Stitt M, Geigenberger P. 2003. ADP-glucose pyrophosphorylase is activated by posttranslational redox-modification in response to light and to sugars in leaves of Arabidopsis and other plant species. Plant Physiol 133: 838–849.

Honjo MN, Emura N, Kawagoe T, Sugisaka J, Kamitani M, Nagano AJ, Kudoh H. 2020. Seasonality of interactions between a plant virus and its host during persistent infection in a natural environment. ISME 14:506–518.

Horton MW, Hancock AM, Huang YS, Toomajian C, Atwell S, Auton A, Muliyati NW, Platt A, Sperone FG, Vilhjalmsson BJ, et al. 2012. Genome-wide patterns of genetic variation in worldwide Arabidopsis thaliana accessions from the RegMap panel. Nat Genet 44: 212–216.

Huot B, Yao J, Montgomery BL, He SY. 2014. Growth–Defense Tradeoffs in Plants: A Balancing Act to Optimize Fitness. Molecular Plant 7: 1267–1287.

Jänkänpää HJ, Mishra Y, SchröDer WP, Jansson S. 2012. Metabolic profiling reveals metabolic shifts in Arabidopsis plants grown under different light conditions: Metabolic profiling under different light regime. Plant, Cell & Environment 35: 1824–1836.

Jenner CE, Walsh JA. 1996. Pathotypic variation in turnip mosaic virus with special reference to European isolates. Plant Pathology 45: 848–856.

Jones RAC. 2016. Future Scenarios for Plant Virus Pathogens as Climate Change Progresses. In: Kielian, M and Maramorosch, K and Mettenleiter T, ed. Advances in Virus Research. Advances in Virus Research. 87–147.

Kafri M, Metzl-Raz E, Jona G, Barkai N. 2015. The Cost of Protein Production. Cell reports 14: 22–31.

Kamitani M, Nagano AJ, Honjo MN, Kudoh H. 2016. RNA-Seq reveals virus–virus and virus–plant interactions in nature. FEMS Microbiology Ecology 92: 1–11.

Kamitani M, Nagano AJ, Honjo MN, Kudoh H. 2018. A survey on plant viruses in natural Brassicaceae community using RNA-Seq. Microb Ecol. 2018. https://doi.org/10.1007/s00248-018-1271-4

Kangasjärvi S, Neukermans J, Li S, Aro E-M, Noctor G. 2012. Photosynthesis, photorespiration, and light signalling in defence responses. Journal of Experimental Botany 63: 1619–1636.

Karasov TL, Chae E, Herman JJ, Bergelson J. 2017. Mechanisms to Mitigate the Trade-Off between Growth and Defense. Plant Cell 29: 666–680.

Kliebenstein DJ. 2004. Secondary metabolites and plant/environment interactions: a view through Arabidopsis thaliana tinged glasses. Plant, Cell & Environment 27: 675–684.

Kliebenstein DJ. 2016. False idolatry of the mythical growth versus immunity tradeoff in molecular systems plant pathology. Physiological and Molecular Plant Pathology 95: 55–59.

Kogovsek P, Pompe-Novak M, Petek M, Fragner L, Weckwerth W, Gruden K. 2016. Primary Metabolism, Phenylpropanoids and Antioxidant Pathways Are Regulated in Potato as a Response to Potato virus Y Infection. PlOS One 11.

Korn M, Gärtner T, Erban A, Kopka J, Selbig J, Hincha DK. 2010. Predicting Arabidopsis freezing tolerance and heterosis in freezing tolerance from metabolite composition. Mol Plant 3: 224–235

Kronheim S, Daniel-Ivad M, Duan Z, Hwang S, Wong AI, Mantel I, Nodwell JR, Maxwell KL. 2018. A chemical defence against phage infection. Nature 564: 283–286.

Likic S, Šola I, Ludwig-Müller J, Rusak G. 2014. Involvement of kaempferol in the defence response of virus infected Arabidopsis thaliana. Eur J Plant Pathol. 138:257–71.

Llave C. 2016. Dynamic cross-talk between host primary metabolism and viruses during infections in plants. Current Opinion in Virology 19: 50–55.

Lozano-Duran R, Zipfel C. 2015. Trade-off between growth and immunity: role of brassinosteroids. Trends in Plant Science 20: 12–19.

Mandadi KK, Scholthof KB. 2013. Plant immune responses against viruses: how does a virus cause disease? Plant Cell 25: 1489–1505.

Melandri G, AbdElgawad H, Riewe D, Hageman JA, Asard H, Beemster GTS, Kadam N, Jagadish K, Altmann T, Ruyter-Spira C, Bouwmeester H. 2019. Biomarkers for grain yield stability in rice under drought stress. Journal of Experimental Botany. doi:10.1093/jxb/erz221

Meyer RC, Steinfath M, Lisec J, Becher M, Witucka-Wall H, Törjék O, Fiehn O, Eckardt A, Willmitzer L, Selbig J., Altmann T. 2007. The metabolic signature related to high plant growth rate in Arabidopsis thaliana. Proc Natl Acad Sci USA 104: 4759–4764

Nagano AJ, Kawagoe T, Sugisaka J, Honjo MN, Iwayama K, Kudoh H. 2019. Annual transcriptome dynamics in natural environments reveals plant seasonal adaptation. Nat Plants. 5:74–83.

Neilson EH, Goodger JQD, Woodrow IE, Moller BL. 2013. Plant chemical defense: at what cost? Trends in Plant Science 18: 250–258.

Nunes-Nesi A, Carrari F, Gibon Y, Sulpice R, Lytovchenko A, Fisahn J, Graham J, Ratcliffe RG, Sweetlove LJ, Fernie AR. 2007. Deficiency of mitochondrial fumarase activity in tomato plants impairs photosynthesis via an effect on stomatal function: Inhibition of fumarase impairs photosynthesis. The Plant Journal 50: 1093–1106.

Obata T, Witt S, Lisec J, Palacios-Rojas N, Florez-Sarasa I, Yousfi S, Araus JL, Cairns JE, Fernie AR. 2015. Metabolite profiles of maize leaves in drought, heat, and combined stress field trials reveal the relationship between metabolism and grain yield. Plant Physiology 169, 2665–2683.

Pagan I, Fraile A, Fernandez-Fueyo E, Montes N, Alonso-Blanco C, Garcia-Arenal F. 2010. Arabidopsis thaliana as a model for the study of plant-virus co-evolution. Philos Trans R Soc Lond B Biol Sci 365: 1983–1995.

Palukaitis P, Yoon J-Y, Choi S-K, Carr JP. 2017. Manipulation of induced resistance to viruses. Current Opinion in Virology 26: 141–148.

Patti GJ, Tautenhahn R, Siuzdak G. 2012. Meta-analysis of untargeted metabolomic data from multiple profiling experiments. Nature Protocols 7: 508–516.

Piasecka A, Jedrzejczak-Rey N, Bednarek P. 2015. Secondary metabolites in plant innate immunity: conserved function of divergent chemicals. New Phytol 206: 948–964.

Pilar Lopez-Gresa M, Lison P, Kim HK, Choi YH, Verpoorte R, Rodrigo I, Conejero V, Maria Belles J. 2012. Metabolic fingerprinting of Tomato Mosaic Virus infected Solanum lycopersicum. Journal of Plant Physiology 169: 1586–1596.

Prasch CM and Sonnewald U. 2013. Simultaneous application of heat, drought, and virus to Arabidopsis plants reveals significant shifts in signaling networks. Plant Physiol. 162: 1849– 1866.

Rasmussen S, Barah P, Suarez-Rodriguez MC, Bressendorff S, Friis P, Costantino P, Bones AM, Nielsen HB, Mundy J. 2013. Transcriptome responses to combinations of stresses in Arabidopsis. Plant Physiol 161: 1783–1794.

Rohart F, Gautier B, Singh A, Le Cao K-A. 2017. mixOmics: an R package for 9omics feature selection and multiple data integration. bioRxiv: 108597.

Rojas CM, Senthil-Kumar M, Tzin V, Mysore KS. 2014. Regulation of primary plant metabolism during plant-pathogen interactions and its contribution to plant defense. Front Plant Sci 5: 17.

Rubio B, Cosson P, Caballero M, Revers F, Bergelson J, Roux F, Schurdi-Levraud V. 2019. Genome wide association study reveals new loci involved in Arabidopsis thaliana and Turnip mosaic virus (TuMV) interactions in the field. New Phytologist 221: 2026–2038.

Sade D, Shriki O, Cuadros-Inostroza A, Tohge T, Semel Y, Haviv Y, Willmitzer L, Fernie AR, Czosnek H, Brotman Y. 2015. Comparative metabolomics and transcriptomics of plant response to Tomato yellow leaf curl virus infection in resistant and susceptible tomato cultivars. Metabolomics 11: 81–97.

Sajitz-Hermstein M, Nikoloski Z. 2010. A novel approach for determining environment-specific protein costs: the case of Arabidopsis thaliana. Bioinformatics 26: i582–8.

Schauer N, Semel Y, Roessner U, Gur A, Balbo I, Carrari F, Pleban T, Perez-Melis A, Bruedigam C, Kopka J, Willmitzer L, Zamir D, Fernie AR. 2006. Comprehensive metabolic profiling and phenotyping of interspecific introgression lines for tomato improvement. Nat. Biotechnol. 24, 447–454.

Shalitin D, Wang Y, Omid A, Gal-On A, Wolf S. 2002. Cucumber mosaic virus movement protein affects sugar metabolism and transport in tobacco and melon plants. Plant Cell Environ 25: 989–997.

Sirikantaramas S, Yamazaki M, Saito K. 2007. Mechanisms of resistance to self-produced toxic secondary metabolites in plants. Phytochemistry Reviews 7: 467.

Smith CA, Want EJ, O’Maille G, Abagyan R, Siuzdak G. 2006. XCMS: Processing Mass Spectrometry Data for Metabolite Profiling Using Nonlinear Peak Alignment, Matching, and Identification. Analytical Chemistry 78: 779–787.

Stare T, Ramšak Ž, Blejec A, Stare K, Turnšek N, Weckwerth W, Wienkoop S, Vodnik D, Gruden K. 2015. Bimodal dynamics of primary metabolism-related responses in tolerant potato-Potato virus Y interaction. BMC Genomics 16.

Steinfath M, Strehmel N, Peters R, Schauer N, Groth D, Hummel J, Steup M, Selbig J, Kopka J, Geigenberger P, van Dongen JT. 2010. Discovering plant metabolic biomarkers for phenotype prediction using an untargeted approach. Plant Biotechnol J 8: 900–911

Stitt M, Quick P. 1989. Photosynthetic carbon partitioning: its regulation and possibilities for manipulation. Physiologia Plantarum 77: 633–641.

Sulpice R, Trenkamp S, Steinfath M, Usadel B, Gibon Y, Witucka-Wall H, Pyl ET, Tschoep H, Steinhauser MC, Guenther M, Hoehne M, Rohwer JM, Altmann T, Fernie AR, Stitt M. 2010. Network analysis of enzyme activities and metabolite levels and their relationship to biomass in a large panel of Arabidopsis accessions. Plant Cell 22: 2872–2893

Sulpice R, Nikoloski Z, Tschoep H, Antonio C, Kleessen S, Larhlimi A, Selbig J, Ishihara H, Gibon Y, Fernie AR, et al. 2013. Impact of the carbon and nitrogen supply on relationships and connectivity between metabolism and biomass in a broad panel of Arabidopsis accessions. Plant Physiol 162: 347–363.

Sulpice R. 2020. Closing the yield gap: can metabolomics be of help? Journal of Experimental Botany 71(2):461–464.

Tecsi LI, Smith AM, Maule AJ, Leegood RC. 1996. A Spatial Analysis of Physiological Changes Associated with Infection of Cotyledons of Marrow Plants with Cucumber Mosaic Virus. Plant Physiol 111: 975–985.

Terce-Laforgue T, Bedu M, Dargel-Grafin C, Dubois F, Gibon Y, Restivo FM, Hirel B. 2013. Resolving the role of plant glutamate dehydrogenase: II. Physiological characterization of plants overexpressing the two enzyme subunits individually or simultaneously. Plant Cell Physiol 54: 1635–1647.

Tian D, Traw MB, Chen JQ, Kreitman M, Bergelson J. 2003. Fitness costs of R-gene-mediated resistance in Arabidopsis thaliana. Nature 423: 74–77

Verma V, Ravindran P, Kumar PP. 2016. Plant hormone-mediated regulation of stress responses. BMC Plant Biology 16.

Wang A. 2015. Dissecting the Molecular Network of Virus-Plant Interactions: The Complex Roles of Host Factors. Annual Review of Phytopathology 53: 45–66.

Wehrens R, Mevik B-H. 2007. The pls package: principal component and partial least squares regression in R.

Younginger BS, Sirová D, Cruzan MB, Ballhorn DJ. 2017. Is biomass a reliable estimate of plant fitness? Applications in Plant Sciences 2017 5(2): 1600094

Zhang N, Gibon Y, Wallace JG, Lepak N, Li P, Dedow L, Chen C, So Y-S, Kremling K, Bradbury PJ, et al. 2015. Genome-Wide association of carbon and nitrogen metabolism in the maize nested association mapping population. Plant Physiology 168: 575–583.

Züst T, Heichinger C, Grossniklaus U, Harrington R, Kliebenstein DJ, Turnbull LA. 2012. Natural enemies drive geographic variation in plant defenses. Science 338: 116–119

