## Supplementary material for "Metabolic profile discriminates and predicts *Arabidopsis* susceptibility to virus under field conditions": Rubio et al. Supplemental information

Text S1: Detailed treatment of metabolite data

·         Removal of peaks present in blanks

·         Retention of peaks present in at least half of QCs

·         Removal of peaks whose QC ‘s variation coefficient is greater than 30 %

·         Correction with the R Package Resource Selection related to the order of the samples

·         Correction based on the dry weight of each sample (in grams)

·         Removal of redundant peaks using clustering methods. Indeed, peaks with masses and/or close retention times could actually correspond to a single peak. Thus, two peaks are considered to be similar if the correlation coefficient is greater than 0.93. In this case, only the peak with the highest intensity is kept.

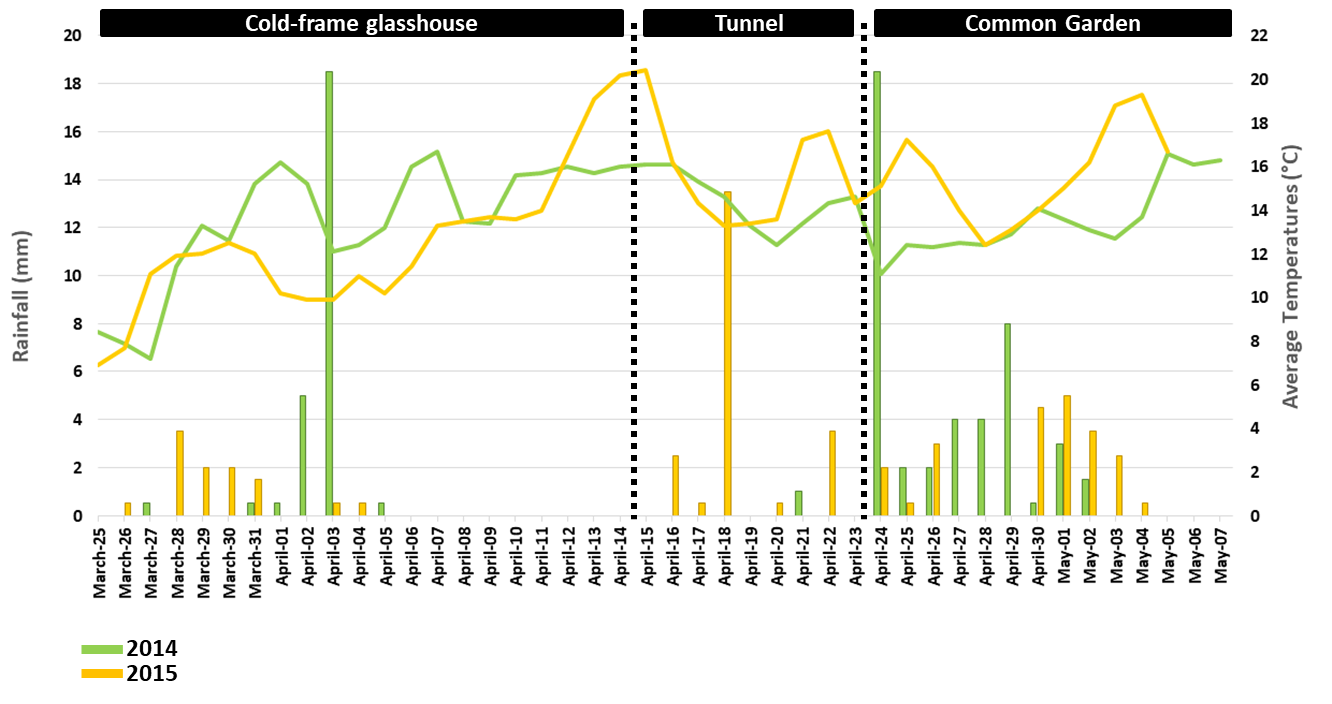

A

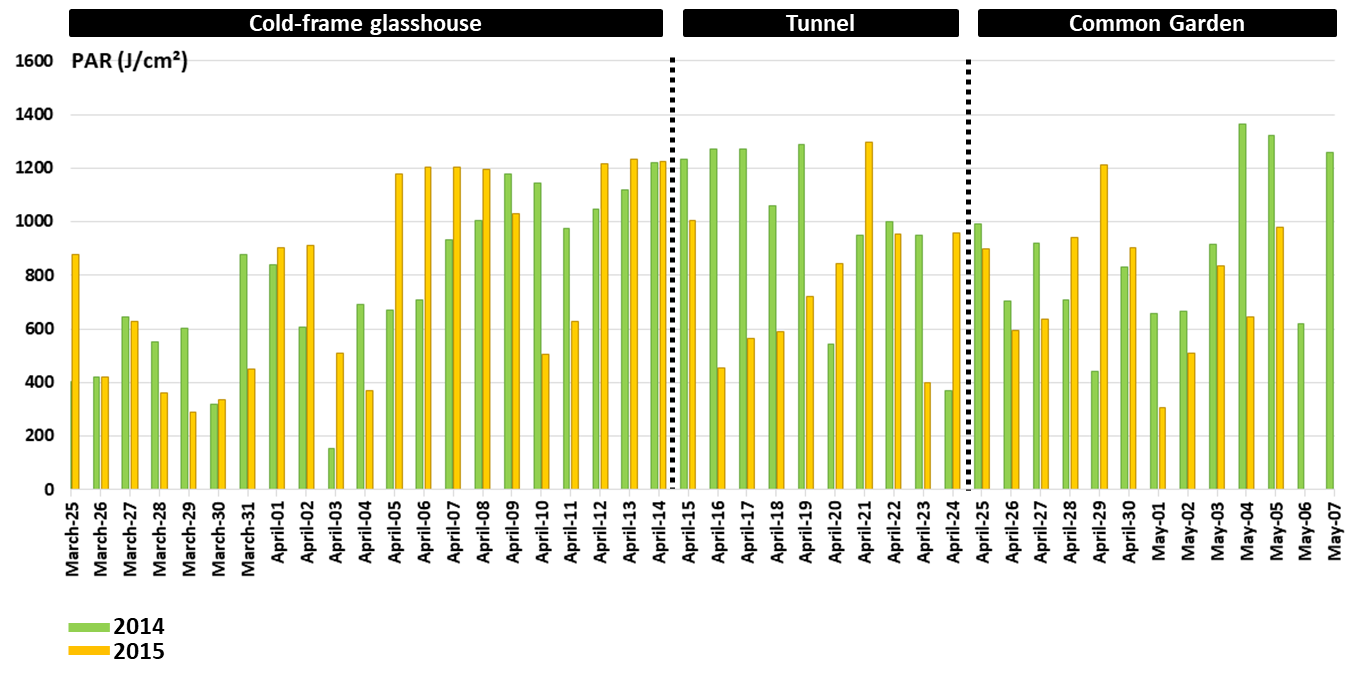

B

C

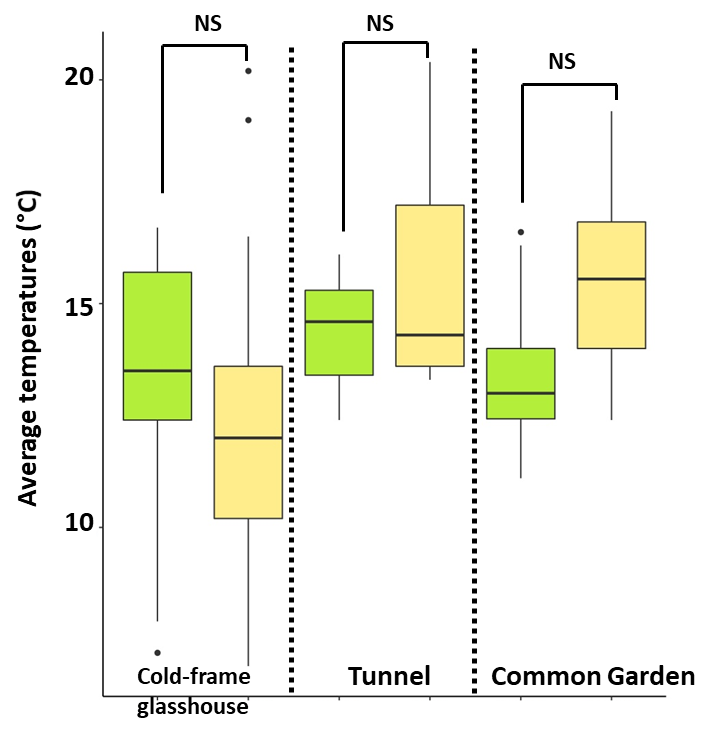

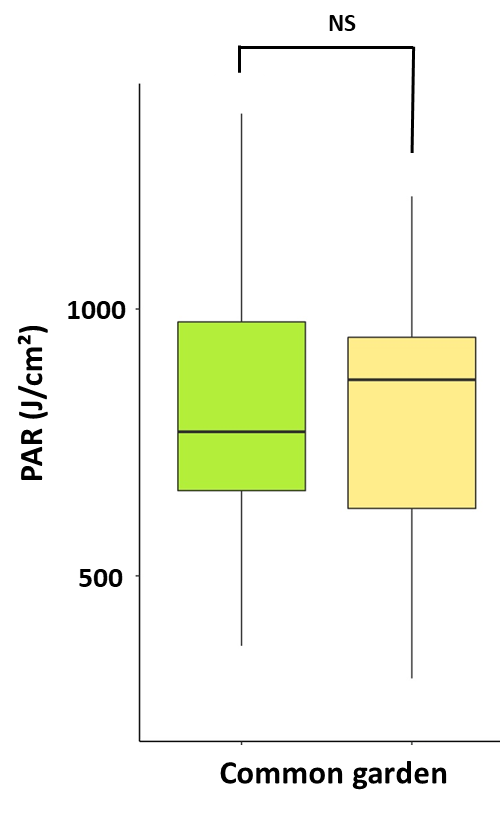

D

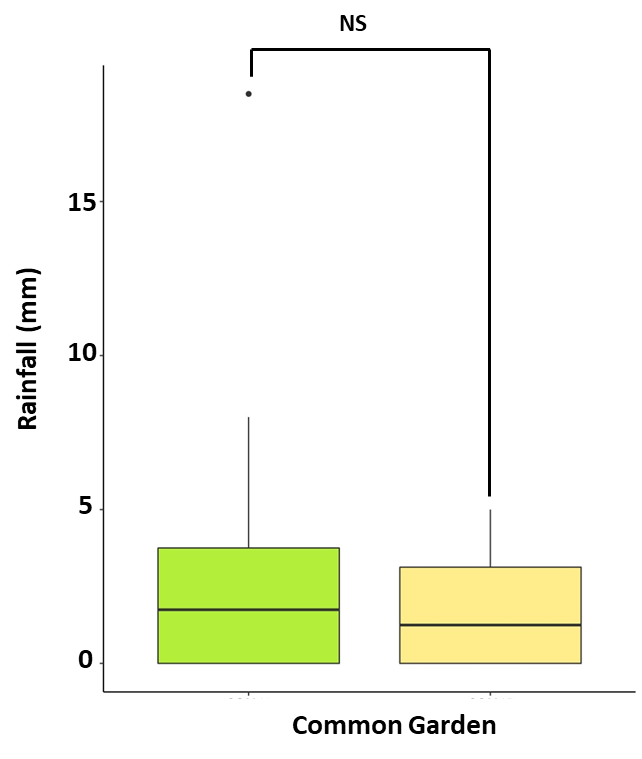

E

**2015**

**2014**

**Figure S1** Climate raw data and comparison between the 2014 and 2015 field experiment

**A**. Rainfall and average temperature in 2014 and 2015 **B**. PAR in 2014 and 2015 recorded under the three successive conditions of the experiment, cold-frame glasshouse, tunnel and common garden. **C**. Comparison of the average temperatures between 2014 and 2015 under the 3 successive conditions of the experiments. **D**. Comparison of rainfall between 2014 and 2015 under common garden conditions. **E**. Comparison of PAR between 2014 and 2015 under common garden conditions. The significance was assessed through a Wilcoxon test at *p-value* = 0.05 indicated by a *, NS: non-significant.

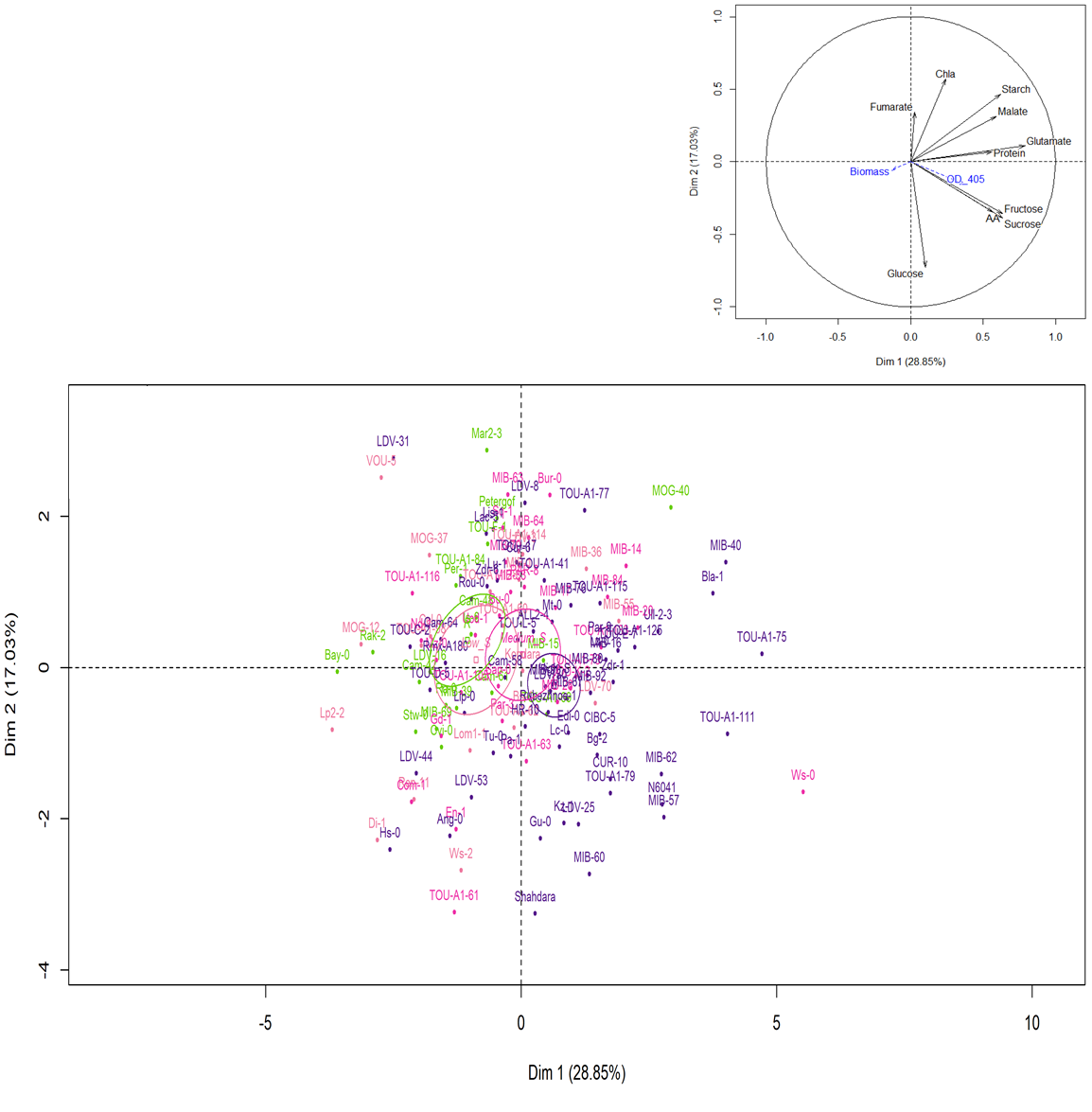

**Figure S2. Score plot of individuals obtained from principal component analysis (PCA) performed on 10 primary metabolic traits measured at 13 days after TuMV inoculation on 130 *A. thaliana* accessions in 2014.**

The two major components that accounted for 45.88% of the variance have been plotted. The confidence ellipses around the centroid of individuals are represented. Categories in color have been defined according to the negative control mock Col-0 which mean OD value was 0.088 (SD 0.01063). In green, infected genotypes with mean OD ≤ 0.088, defined as resistant. In light pink, infected genotypes with 0.088<mean OD≤2*0.088. In pink, infected genotypes with 2*0.088<mean OD≤3*0.088. In purple, infected genotypes with mean OD>3*0.088. All these three categories were defined as susceptible as described in table S1.

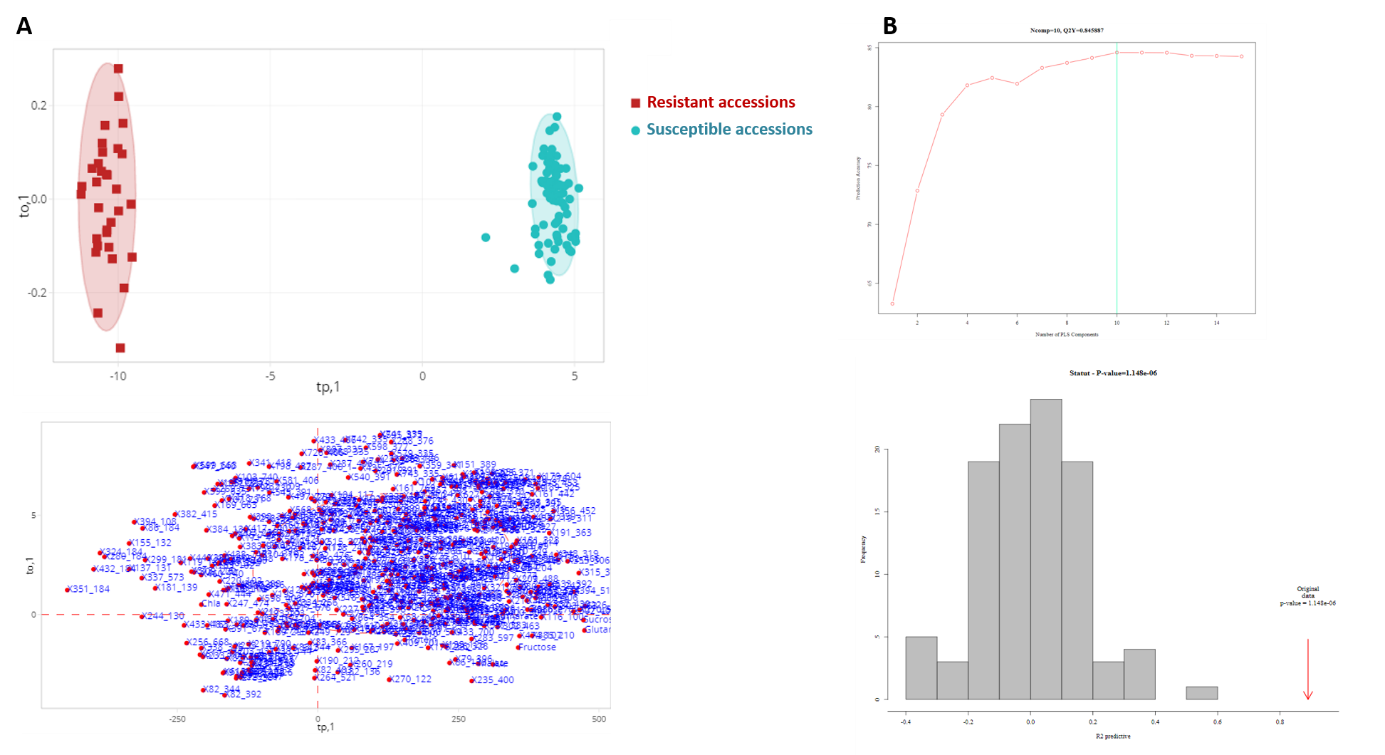

**Figure S3** **OPLS-DA analysis and its parameters of validation for the TuMV-inoculated resistant and susceptible 26 *A. thaliana* accessions of the ‘2015’ field experiment A.** OPLS-DA results with the score plot and the loading plot containing all the metabolic variables tested. The resistant (in red) and susceptible (in blue) accessions were represented. **B.** Parameters for validation of the OPLS-DA analysis.

**Table S1** **List of the A. thaliana accessions with their geographic position and their susceptible status to TuMV infection confirmed by OD values and SD in both experiments**

| **Genotype** | **ID** | **Latitude** | **Longitude** | **Country** | **Susceptibility Groups^1^** | **OD Means** | | **OD-Standard Deviations** | |
| --- | --- | --- | --- | --- | --- | --- | --- | --- | --- |
|  |  |  |  |  |  | **2014** | **2015** | **2014** | **2015** |
| Bay-0 | 6899 | 49 | 11 | GER | **R** | **0.009** | 0.004 | 0.0138 | 0.0155 |
| Cam-42 | 45 | 48.2667 | -4.58333 | FRA | **R** | **0.023** | - | 0.0149 | - |
| Cam-48 | 51 | 48.2667 | -4.58333 | FRA | **R** | **-0.001** | - | 0.0148 | - |
| Cam-61 | 66 | 48.2667 | -4.58333 | FRA | **R** | **0.004** | - | 0.0035 | - |
| Cvi-0 | 6911 | 15.1111 | -23.6167 | CPV | **R** | **-0.009** | - | 0.0183 | - |
| Is-0 | 8312 | 50.5 | 7.5 | GER | **R** | **0.016** | - | 0.0191 | - |
| LDV-16 | 106 | 48.5167 | -4.06667 | FRA | **R** | **0.011** | - | 0.0015 | - |
| Mar2-3 | 159 | 47.35 | 3.93333 | FRA | **R** | **0.002** | 0.005 | 0.0024 | 0.0087 |
| MIB-15 | 166 | 47.3833 | 5.31667 | FRA | **R** | **0.002** | - | 0.0021 | - |
| MIB-39 | 190 | 47.3833 | 5.31667 | FRA | **R** | **0.011** | 0.007 | 0.0055 | 0.0155 |
| MIB-69 | 212 | 47.3833 | 5.31667 | FRA | **R** | **0.020** | - | 0.0175 | - |
| MOG-40 | 244 | 48.6667 | -4.06667 | FRA | **R** | **0.017** | - | 0.0163 | - |
| Per-1 | 8354 | 58 | 56.3167 | RUS | **R** | **0.037** | - | 0.056 | - |
| Petergof | 7296 | 59 | 29 | RUS | **R** | **0.006** | 0.011 | 0.0045 | 0.0252 |
| Ra-0 | 6958 | 46 | 3.3 | FRA | **R** | **0.024** | 0.003 | 0.0156 | 0.0072 |
| Rak-2 | 8365 | 49 | 16 | CZE | **R** | **0.004** | 0.006 | 0.0052 | 0.00204 |
| Stw-0 | 8388 | 52 | 36 | RUS | **R** | **0.029** | - | 0.0369 | - |
| TOU-A1-69 | 335 | 46.6667 | 4.11667 | FRA | **R** | **-0.002** | 0.015 | 0.0109 | 0.0224 |
| TOU-A1-84 | 348 | 46.6667 | 4.11667 | FRA | **R** | **0.028** | 0.007 | 0.022 | 0.0204 |
| TOU-F-1 | 369 | 46.6667 | 4.11667 | FRA | **R** | **0.023** | - | 0.0135 | - |
| Ws-2 | 6981 | 52.3 | 30 | RUS | **S** | **0.092** | - | - | - |
| TOU-A1-98 | 359 | 46.6667 | 4.11667 | FRA | **S** | **0.101** | - | - | - |
| TOU-A1-114 | 280 | 46.6667 | 4.11667 | FRA | **S** | **0.101** | - | - | - |
| MOG-12 | 237 | 48.6667 | -4.06667 | FRA | **S** | **0.105** | - | - | - |
| TOU-A1-62 | 328 | 46.6667 | 4.11667 | FRA | **S** | **0.108** | - | - | - |
| Kondara | 6929 | 38.48 | 68.49 | TJK | **S** | **0.110** | - | - | - |
| Lp2-2 | 7520 | 49.38 | 16.81 | CZE | **S** | **0.119** | - | - | - |
| MOG-37 | 242 | 48.6667 | -4.06667 | FRA | **S** | **0.124** | - | - | - |
| MIB-36 | 187 | 47.3833 | 5.31667 | FRA | **S** | **0.125** | - | 0.0127 | - |
| TOU-A1-60 | 326 | 46.6667 | 4.11667 | FRA | **S** | **0.131** | - | - | - |
| MIB-55 | 201 | 47.3833 | 5.31667 | FRA | **S** | **0.134** | - | 0.0477 | - |
| MIB-1 | 160 | 47.3833 | 5.31667 | FRA | **S** | **0.138** | - | 0.0191 | - |
| TOU-A1-128 | 292 | 46.6667 | 4.11667 | FRA | **S** | **0.140** | - | - | - |
| LDV-3 | 121 | 48.5167 | -4.06667 | FRA | **S** | **0.140** | - | 0.0507 | - |
| Bs-1 | 8270 | 47.5 | 7.5 | SUI | **S** | **0.147** | 0.66 | 0.0696 | 0.339 |
| VOU-5 | 394 | 46.65 | 0.166667 | FRA | **S** | **0.154** | - | - | - |
| Lom-1-1 | 6042 | 56.09 | 13.9 | SWE | **S** | **0.155** | - | - | - |
| LDV-70 | 156 | 48.5167 | -4.06667 | FRA | **S** | **0.165** | - | - | - |
| Col-0 | 6909 | 38.3 | -92.3 | USA | **S** | **0.165** | 0.391 | 0.0839 | 0.215 |
| Ren-11 | 6960 | 48.5 | -1.41 | FRA | **S** | **0.172** | - | - | - |
| Di-1 | 7098 | 47 | 5 | FRA | **S** | **0.175** | - | 0.0012 | - |
| MIB-63 | 207 | 47.3833 | 5.31667 | FRA | **S** | **0.182** | - | 0.1158 | - |
| Gd-1 | 8296 | 53.5 | 10.5 | GER | **S** | **0.186** | - | 0.069 | - |
| Mz-0 | 6940 | 50.3 | 8.3 | GER | **S** | **0.190** | 0.59 | 0.0189 | 0.263 |
| TOU-K-2 | 385 | 46.6667 | 4.11667 | FRA | **S** | **0.194** | - | 0.0339 | - |
| MIB-33 | 184 | 47.3833 | 5.31667 | FRA | **S** | **0.197** | - | 0.0731 | - |
| MIB-64 | 208 | 47.3833 | 5.31667 | FRA | **S** | **0.198** | - | 0.0721 | - |
| PAR-8 | 262 | 46.65 | -0.25 | FRA | **S** | **0.198** | - | 0.0876 | - |
| Bu-0 | 8271 | 50.5 | 9.5 | GER | **S** | **0.201** | 0.4 | - | 0.203 |
| MIB-20 | 171 | 47.3833 | 5.31667 | FRA | **S** | **0.202** | 0.525 | 0.0712 | 0.294 |
| Par-3 | 258 | 46.65 | -0.25 | FRA | **S** | **0.202** | - | 0.0878 | - |
| TOU-A1-73 | 338 | 46.6667 | 4.11667 | FRA | **S** | **0.205** | 0.55 | 0.1267 | 0.261 |
| En-1 | 8290 | 50 | 8.5 | GER | **S** | **0.210** | - | - | - |
| MIB-28 | 178 | 47.3833 | 5.31667 | FRA | **S** | **0.211** | 0.428 | 0.073 | 0.242 |
| TOU-A1-116 | 282 | 46.6667 | 4.11667 | FRA | **S** | **0.219** | - | 0.1391 | - |
| TOU-A1-61 | 327 | 46.6667 | 4.11667 | FRA | **S** | **0.219** | - | 0.0418 | - |
| Com-1 | 7092 | 49.416 | 2.823 | FRA | **S** | **0.220** | - | 0.0314 | - |
| Sq-1 | 6966 | 51.4083 | -0.6383 | UK | **S** | **0.220** | - | - | - |
| MIB-17 | 168 | 47.3833 | 5.31667 | FRA | **S** | **0.227** | - | - | - |
| TOU-A1-133 | 296 | 46.6667 | 4.11667 | FRA | **S** | **0.228** | - | 0.0582 | - |
| Nd-1 | 6942 | 50 | 10 | SUI | **S** | **0.229** | - | 0.1933 | - |
| TOU-A1-105 | 273 | 46.6667 | 4.11667 | FRA | **S** | **0.233** | - | 0.0428 | - |
| MIB-84 | 223 | 47.3833 | 5.31667 | FRA | **S** | **0.234** | - | 0.0285 | - |
| Bur-0 | 5719 | 54.1 | -6.2 | IRL | **S** | **0.239** | - | - | - |
| MIB-70 | 213 | 47.3833 | 5.31667 | FRA | **S** | **0.244** | - | - | - |
| TOU-A1-63 | 329 | 46.6667 | 4.11667 | FRA | **S** | **0.249** | - | 0.1233 | - |
| Sap-0 | 8378 | 49.49 | 14.24 | CZE | **S** | **0.250** | - | 0.1215 | - |
| Uod-1 | 6975 | 48.3 | 14.45 | AUT | **S** | **0.260** | - | - | - |
| MIB-14 | 165 | 47.3833 | 5.31667 | FRA | **S** | **0.265** | - | - | - |
| Ws-0 | 6980 | 52.3 | 30 | RUS | **S** | **0.265** | - | - | - |
| Hs-0 | 8310 | 52.24 | 9.44 | GER | **S** | **0.274** | 0.46 | 0.1605 | 0.212 |
| TOU-A1-111 | 277 | 46.6667 | 4.11667 | FRA | **S** | **0.276** | - | 0.1237 | - |
| MIB-57 | 202 | 47.3833 | 5.31667 | FRA | **S** | **0.284** | - | 0.1119 | - |
| CIBC-5 | 6730 | 51.4083 | -0.6383 | UK | **S** | **0.287** | - | - | - |
| MIB-80 | 219 | 47.3833 | 5.31667 | FRA | **S** | **0.292** | - | - | - |
| Ull-2-3 | 6973 | 56.0648 | 13.9707 | SWE | **S** | **0.292** | - | - | - |
| Lc-0 | 8323 | 57 | -4 | UK | **S** | **0.296** | - | - | - |
| Gu-0 | 6922 | 50.3 | 8 | GER | **S** | **0.298** | 0.389 | 0.1019 | 0.272 |
| MIB-16 | 167 | 47.3833 | 5.31667 | FRA | **S** | **0.298** | - | - | - |
| TOU-A1-75 | 340 | 46.6667 | 4.11667 | FRA | **S** | **0.299** | - | 0.1969 | - |
| LDV-53 | 146 | 48.5167 | -4.06667 | FRA | **S** | **0.300** | - | 0.204 | - |
| Rubezhnoe-1 | 7323 | 49 | 38.28 | UKR | **S** | **0.300** | - | - | - |
| MIB-92 | 230 | 47.3833 | 5.31667 | FRA | **S** | **0.302** | - | 0.1252 | - |
| TOU-I-17 | 378 | 46.6667 | 4.11667 | FRA | **S** | **0.302** | - | - | - |
| LDV-31 | 123 | 48.5167 | -4.06667 | FRA | **S** | **0.306** | - | - | - |
| LDV-8 | 157 | 48.5167 | -4.06667 | FRA | **S** | **0.308** | - | - | - |
| Tu-0 | 8395 | 45 | 7.5 | ITA | **S** | **0.312** | - | 0.0395 | - |
| Ang-0 | 6992 | 50.3 | 5.3 | BEL | **S** | **0.316** | - | - | - |
| LDV-44 | 137 | 48.5167 | -4.06667 | FRA | **S** | **0.318** | - | - | - |
| TOU-C-2 | 361 | 46.6667 | 4.11667 | FRA | **S** | **0.319** | - | - | - |
| MIB-90 | 229 | 47.3833 | 5.31667 | FRA | **S** | **0.325** | - | 0.2399 | - |
| TOU-A1-77 | 341 | 46.6667 | 4.11667 | FRA | **S** | **0.328** | - | - | - |
| TOU-D-5 | 364 | 46.6667 | 4.11667 | FRA | **S** | **0.329** | - | - | - |
| Lip-0 | 8325 | 50 | 19.3 | POL | **S** | **0.330** | - | - | - |
| Rmx-A180 | 7525 | 42.036 | -86.511 | USA | **S** | **0.339** | - | - | - |
| Lz-0 | 6936 | 46 | 3.3 | FRA | **S** | **0.341** | - | - | - |
| Lu-1 | 8334 | 55.71 | 13.2 | SWE | **S** | **0.341** | - | - | - |
| HR-10 | 6923 | 51.4083 | -0.6383 | UK | **S** | **0.343** | - | 0.0332 | - |
| Cur-6 | 84 | 45 | 1.75 | FRA | **S** | **0.352** | - | 0.2236 | - |
| Lac-5 | 96 | 47.7 | 6.81667 | FRA | **S** | **0.368** | - | - | - |
| Zdr-1 | 6984 | 49.3853 | 16.2544 | CZE | **S** | **0.369** | - | 0.1718 | - |
| Kz-1 | 6930 | 49.5 | 73.1 | KAZ | **S** | **0.377** | - | 0.1481 | - |
| Bla-1 | 7015 | 41.6833 | 2.8 | ESP | **S** | **0.381** | - | - | - |
| Cam-64 | 69 | 48.2667 | -4.58333 | FRA | **S** | **0.407** | - | - | - |
| Pa-1 | 8353 | 38.07 | 13.22 | ITA | **S** | **0.409** | - | 0.0787 | - |
| MIB-76 | 216 | 47.3833 | 5.31667 | FRA | **S** | **0.411** | - | 0.0161 | - |
| TOU-E-7 | 368 | 46.6667 | 4.11667 | FRA | **S** | **0.414** | - | 0.0825 | - |
| TOU-A1-125 | 291 | 46.6667 | 4.11667 | FRA | **S** | **0.418** | 0.403 | - | 0.147 |
| Par-9 | 263 | 46.65 | -0.25 | FRA | **S** | **0.429** | - | - | - |
| MIB-67 | 210 | 47.3833 | 5.31667 | FRA | **S** | **0.434** | 0.549 | 0.1683 | 0.249 |
| TOU-A1-79 | 343 | 46.6667 | 4.11667 | FRA | **S** | **0.436** | - | - | - |
| Edi-0 | 6914 | 56 | -3 | UK | **S** | **0.438** | 0.258 | 0.086 | 0.169 |
| Bg-2 | 6709 | 47.6479 | -122.305 | USA | **S** | **0.441** | 0.324 | - | 0.211 |
| Mt-0 | 6939 | 32.34 | 22.46 | LIB | **S** | **0.450** | 0.467 | 0.0387 | 0.2 |
| TOU-A1-41 | 320 | 46.6667 | 4.11667 | FRA | **S** | **0.451** | - | 0.2366 | - |
| ALL2-4 | 12 | 45.2667 | 1.48333 | FRA | **S** | **0.462** | - | 0.2833 | - |
| CUR-10 | 79 | 45 | 1.75 | FRA | **S** | **0.463** | 0.576 | - | 0.147 |
| Shahdara | 6962 | 38.35 | 68.48 | TJK | **S** | **0.467** | - | 0.0453 | - |
| MIB-40 | 191 | 47.3833 | 5.31667 | FRA | **S** | **0.474** | - | - | - |
| LDV-30 | 122 | 48.5167 | -4.06667 | FRA | **S** | **0.492** | - | 0.2742 | - |
| Cam-58 | 62 | 48.2667 | -4.58333 | FRA | **S** | **0.496** | - | 0.1592 | - |
| MIB-62 | 206 | 47.3833 | 5.31667 | FRA | **S** | **0.507** | 0.346 | - | 0.2084 |
| N6041 | 8420 | 50.0667 | 8.5333 | GER | **S** | **0.521** | - | - | - |
| TOU-L-5 | 389 | 46.6667 | 4.11667 | FRA | **S** | **0.570** | 0.407 | 0.5299 | 0.19 |
| Zdr-6 | 6985 | 49.3853 | 16.2544 | CZE | **S** | **0.578** | - | - | - |
| Rou-0 | 7320 | 49.4424 | 1.09849 | FRA | **S** | **0.611** | - | 0.5242 | - |
| LDV-25 | 116 | 48.5167 | -4.06667 | FRA | **S** | **0.622** | - | 0.2129 | - |
| Lis-1 | 8326 | 56 | 14.7 | SWE | **S** | **0.679** | - | - | - |
| TOU-A1-115 | 281 | 46.6667 | 4.11667 | FRA | **S** | **0.750** | - | - | - |
| MIB-60 | 204 | 47.3833 | 5.31667 | FRA | **S** | **0.896** | 0.425 | - | 0.145 |

^1^ In 2014, categories in color have been defined according to the healthy control Col-0 which mean OD value was 0.088 (SD 0.01063).

In green, infected genotypes with mean OD ≤ 0.088, defined as resistant (R).

In light pink to purple, accessions are classified as susceptible (S). In light pink, infected genotypes with 0.088<mean OD≤2*0.088. In pink, infected genotypes with 2*0.088<mean OD≤3*0.088. In purple, infected genotypes with mean OD>3*0.088. All these three categories were defined as susceptible.

**Table S2 Spearman correlations between dry biomass, viral accumulation (OD) and primary metabolites content measured (A) on the 130 *A. thaliana* accessions of the ‘2014’ field experiment and (B) on the 26 *A. thaliana* accessions of the ‘2015’ field experiment**

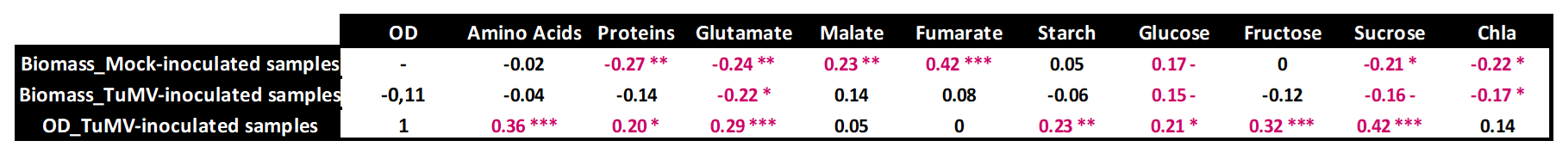

A

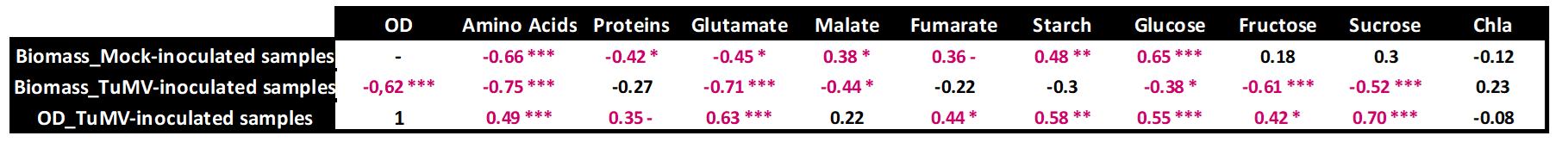

B

All the significant correlations where showed by asterisks and values (Signif.codes: 0 ‘***’ 0.001 ‘**’ 0.01 ‘*’ 0.05 ‘.’ 0.1)

- Non tested

**Table S3** Common Variable Importance in the Projection (VIPs) obtained with the OPLS-DA performed on mock and TuMV-inoculated and on resistant and susceptible accessions of twenty-six accessions in 2015. For each VIP, the comparison between the metabolic content in mock (M) and TuMV-inoculated (I) samples was done and the fold change was represented. Moreover, the comparison, for each VIP, of metabolic content between resistant (R) and susceptible (S) accessions was also represented.

| **VIP-OPLS-DA^1^** | **VIP values** | **m/z^2^** | **rt^3^** | **Mock vs. TuMV Infected** | **Fold change** | **Resistant vs. Susceptible** |
| --- | --- | --- | --- | --- | --- | --- |
| **193;324** | **2.201000988** | 193.13 | 323.817 | **I > M** | **4.54** | **S > R ***^4^** |
| **331;329** | **2.177409793** | 331.12 | 328.817 | **I > M** | **2.55** | **S > R ***** |
| **474;107** | **2.100348101** | 474.22 | 107.298 | **I > M** | **7.12** | **S > R ***** |
| **482;101** | **2.02851797** | 482.11 | 101.274 | **I > M** | **3.24** | **S > R ***** |
| **205;242** | **1.95494017** | 205.1 | 242.121 | **I > M** | **3.16** | **S > R ***** |
| **209;488** | **1.895244441** | 209.15 | 488.35 | **I > M** | **3.87** | **S > R ***** |
| **212;284** | **1.812272811** | 211.56 | 283.891 | **I > M** | **1.97** | **S > R ***** |
| **343;345** | **1.793276838** | 343.12 | 345.319 | **I > M** | **2.41** | **S > R ***** |
| **116;100** | **1.77949346** | 116.07 | 100.163 | **I > M** | **6.88** | **S > R ***** |
| **162;357** | **1.767230127** | 162.05 | 356.879 | **I > M** | **3.1** | **S > R ***** |
| **303;463** | **1.682969043** | 303.13 | 463.27 | **I > M** | **2.9** | **S > R ***** |
| **219;311** | **1.640982077** | 219.1 | 311.134 | **I > M** | **3.44** | **S > R ***** |
| **355;306** | **1.594996482** | 355.1 | 306.288 | **I > M** | **8.11** | **S > R ***** |
| **305;210** | **1.575910966** | 305.09 | 210.166 | **I > M** | **2.69** | **S > R ***** |
| **133;296** | **1.573543164** | 133.06 | 295.68 | **I > M** | **3.79** | **S > R ***** |
| **221;216** | **1.49354017** | 221.09 | 215.744 | **I > M** | **2.46** | **S > R ***** |
| **124;346** | **1.493002053** | 124.08 | 346.354 | **I > M** | **1.68** | **S > R ***** |
| **503;391** | **1.478656897** | 503.19 | 391.457 | **I > M** | **2.24** | **S > R ***** |
| **373;344** | **1.456529478** | 373.13 | 343.531 | **I > M** | **1.78** | **S > R ***** |
| **151;359** | **1.45485619** | 151.08 | 359.377 | **I > M** | **1.28** | **S > R ***** |
| **374;322** | **1.444521384** | 374.14 | 321.703 | **I > M** | **1.88** | **S > R ***** |
| **162;249** | **1.431791986** | 162.05 | 249.101 | **I > M** | **1.81** | **S > R ***** |
| **175;402** | **1.423091039** | 175.15 | 401.831 | **I > M** | **2.52** | **S > R ***** |
| **332;430** | **1.416086097** | 332.13 | 430.481 | **I > M** | **2.32** | **S > R ***** |
| **315;448** | **1.412434025** | 315.13 | 447.821 | **I > M** | **2.05** | **S > R ***** |
| **221;230** | **1.4121623** | 221.12 | 230.244 | **I > M** | **1.67** | **S > R ***** |
| **391;774** | **1.397491479** | 391.24 | 774.477 | **I > M** | **2.36** | **S > R ***** |
| **386;278** | **1.385282143** | 386.22 | 277.549 | **I > M** | **5.51** | **S > R ***** |
| **348;319** | **1.376257143** | 348.27 | 318.945 | **I > M** | **4.2** | **S > R ***** |
| **356;452** | **1.358448769** | 356.12 | 452.352 | **I > M** | **1.69** | **S > R ***** |
| **201;344** | **1.355039751** | 201.05 | 343.832 | **I > M** | **1.71** | **S > R ***** |
| **222;216** | **1.351138929** | 221.6 | 215.822 | **I > M** | **2.46** | **S > R **** |
| **370;302** | **1.331366534** | 370.15 | 301.519 | **I > M** | **5.59** | **S > R ***** |
| **315;370** | **1.330904866** | 315.13 | 370.476 | **I > M** | **3.58** | **S > R ***** |
| **367;358** | **1.251165644** | 367.1 | 358.491 | **I > M** | **1.62** | **S > R ***** |
| **182;466** | **1.243641489** | 182.08 | 465.796 | **I > M** | **3.65** | **S > R ***** |
| **302;407** | **1.221943812** | 302.1 | 406.697 | **I > M** | **4.3** | **S > R ***** |
| **191;363** | **1.219391802** | 191.07 | 362.643 | **I > M** | **2.37** | **S > R ***** |
| **367;342** | **1.199530791** | 367.15 | 341.908 | **I > M** | **2.4** | **S > R ***** |
| **291;259** | **1.165429681** | 291.18 | 259.063 | **I > M** | **3.68** | **S > R ***** |
| **178;191** | **1.128214863** | 178.09 | 191.013 | **I > M** | **3.11** | **S > R ***** |
| **396;348** | **1.126209569** | 396.11 | 347.739 | **I > M** | **7.35** | **S > R ***** |
| **394;517** | **1.125664243** | 394.2 | 517.072 | **I > M** | **4.36** | **S > R ***** |
| **543;99** | **1.122326073** | 543.13 | 98.5221 | **I > M** | **2.62** | **S > R ***** |
| **212;774** | **1.113329112** | 212.09 | 774.047 | **I > M** | **3.14** | **S > R ***** |
| **164;375** | **1.107007943** | 164.07 | 375.012 | **I > M** | **2.38** | **S > R ***** |
| **210;465** | **1.09694612** | 210.11 | 464.799 | **I > M** | **2.27** | **S > R ***** |
| **379;402** | **1.091061829** | 379.09 | 402.311 | **I > M** | **2.47** | **S > R ***** |
| **133;126** | **1.06741008** | 133.1 | 126.347 | **I > M** | **2.62** | **S > R **** |
| **409;491** | **1.066982099** | 409.17 | 490.545 | **I > M** | **1.46** | **S > R ***** |
| **109;359** | **1.035812334** | 109.06 | 359.34 | **I > M** | **1.67** | **S > R ***** |
| **162;402** | **1.032043124** | 162.05 | 402.281 | **I > M** | **1.41** | **S > R ***** |
| **227;789** | **1.023548533** | 227.16 | 788.506 | **I > M** | **1.71** | **S > R ***** |
| **192;462** | **1.000060126** | 192.04 | 462.37 | **I > M** | **1.31** | **S > R ***** |
| **433;700** | **2.253892757** | 433.24 | 700.261 | **I > M** | **3** | **R > S ***** |
| **361;491** | **1.914433317** | 361.09 | 490.564 | **I > M** | **4.84** | **R > S ***** |
| **512;491** | **1.813252344** | 512.13 | 491.367 | **I > M** | **21.08** | **R > S ***** |
| **64;395** | **1.668700459** | 63.934 | 395.047 | **I > M** | **1.25** | **R > S ***** |
| **79;396** | **2.194133112** | 79.041 | 395.754 | **M > I** | **1.85** | **S > R ***** |
| **105;700** | **1.816365199** | 105.07 | 700.287 | **M > I** | **1.88** | **S > R ***** |
| **169;496** | **1.288454553** | 169.05 | 495.5 | **M > I** | **1.91** | **S > R ***** |
| **449;442** | **1.102028381** | 449.11 | 442.311 | **M > I** | **4.26** | **R > S ***** |
| **137;131** | **1.036410357** | 136.93 | 131.31 | **M > I** | **1.35** | **R > S ***** |

^1^ When undetermined, VIP are identified through m/z;rt values. VIP values are classified in decreasing order.

^2^ mass to charge ratio

^3^ retention time

^4^ The significance was assessed through a Wilcoxon test at *** *P* < 0.001, ** 0.001 < P < 0.01

**Table S4** Comparisons between the Variable Importance in the Projection (VIPs) identify by OPLS-DA analysis and those identify by PLS analysis performed with resistant and susceptible twenty-six accessions in 2015. The same results of metabolic content comparisons were found between resistant (R) and susceptible (S) accessions. The fold change was also represented for each VIP. Primary metabolites are light-grey highlighted. Metabolites that accumulate significantly more in resistant accessions are at the bottom of the table.

| **VIP PLS^1^** | **VIP-PLS value** | **VIP-OPLS-DA value** | **m/z^2^** | **rt^3^** | **Resistant vs Susceptible metabolic contents** | **Fold change** |
| --- | --- | --- | --- | --- | --- | --- |
| 356;452 | 2.1040 | 1.9580 | 356.1202 | 452.352 | S > R ***^4^ | 2.17 |
| Glutamate | 2.0854 | 1.5902 | NA | NA | S > R *** | 1.47 |
| Sucrose | 1.9944 | 2.2126 | NA | NA | S > R *** | 2.89 |
| 219;311 | 1.9753 | 1.9395 | 219.1011 | 311.134 | S > R *** | 4.81 |
| 385;210 | 1.9524 | 1.7380 | 385.1055 | 210.083 | S > R *** | 45.77 |
| 373;453 | 1.9059 | 1.2900 | 373.1273 | 452.6 | S > R *** | 1.81 |
| 270;587 | 1.8532 | 1.5343 | 270.133 | 587.091 | S > R *** | 43.68 |
| 315;370 | 1.8277 | 1.5236 | 315.1333 | 370.476 | S > R *** | 3.93 |
| 348;319 | 1.8190 | 1.3963 | 348.2736 | 318.945 | S > R *** | 3.61 |
| 394;517 | 1.8096 | 1.4897 | 394.2045 | 517.072 | S > R *** | 17.28 |
| 191;363 | 1.7958 | 1.3599 | 191.0699 | 362.643 | S > R *** | 2.76 |
| 474;107 | 1.7890 | 1.6748 | 474.2178 | 107.298 | S > R *** | 2.68 |
| 226;406 | 1.7712 | 1.5540 | 226.1066 | 406.369 | S > R *** | 7.97 |
| 420;276 | 1.7295 | 1.2167 | 419.6949 | 275.797 | S > R *** | 1.77 |
| 367;342 | 1.7210 | 1.0107 | 367.1531 | 341.908 | S > R *** | 1.96 |
| 302;407 | 1.6916 | 1.4864 | 302.1013 | 406.697 | S > R *** | 7.16 |
| 503;391 | 1.6879 | 1.6947 | 503.1902 | 391.457 | S > R *** | 2.51 |
| 116;100 | 1.6790 | 1.8669 | 116.0701 | 100.163 | S > R *** | 6.05 |
| 396;257 | 1.6725 | 1.1064 | 396.1854 | 256.51 | S > R *** | 1.43 |
| 303;463 | 1.6577 | 1.9734 | 303.1334 | 463.27 | S > R *** | 3.69 |
| 355;306 | 1.6414 | 1.4506 | 355.1015 | 306.288 | S > R *** | 3.29 |
| 370;302 | 1.6342 | 1.6097 | 370.1486 | 301.519 | S > R *** | 28.49 |
| 343;345 | 1.6096 | 1.6930 | 343.1168 | 345.319 | S > R *** | 2.08 |
| 533;392 | 1.6000 | 1.9258 | 533.1549 | 391.765 | S > R *** | 4.98 |
| 201;451 | 1.5993 | 1.0635 | 201.0543 | 451.123 | S > R *** | 1.75 |
| 757;319 | 1.5938 | 1.5374 | 757.2171 | 319.13 | S > R *** | 1.69 |
| 386;278 | 1.5899 | 1.2560 | 386.2198 | 277.549 | S > R *** | 3.48 |
| 903;319 | 1.5874 | 1.7102 | 903.2769 | 318.889 | S > R *** | 8.48 |
| 449;319 | 1.5823 | 1.5451 | 449.1063 | 318.949 | S > R *** | 1.75 |
| 182;466 | 1.5645 | 1.4146 | 182.0808 | 465.796 | S > R *** | 4.21 |
| 175;402 | 1.5414 | 1.7128 | 175.1478 | 401.831 | S > R *** | 3.77 |
| 305;210 | 1.5354 | 1.4081 | 305.0861 | 210.166 | S > R *** | 2.24 |
| 195;453 | 1.5352 | 1.1541 | 195.0648 | 452.614 | S > R *** | 1.55 |
| 221;216 | 1.5350 | 1.7483 | 221.0915 | 215.744 | S > R *** | 3.13 |
| 374;322 | 1.5256 | 1.3048 | 374.1436 | 321.703 | S > R *** | 1.77 |
| Glucose | 1.5212 | 1.7342 | NA | NA | S > R *** | 2.49 |
| 179;604 | 1.5211 | 1.3057 | 179.1063 | 603.961 | S > R *** | 1.64 |
| 161;442 | 1.5180 | 1.2744 | 161.0957 | 441.797 | S > R *** | 1.79 |
| **394;108** | 1.7935 | **1.5477** | **394.2001** | **108.419** | **R > S ***** | **1.93** |
| **351;184** | 1.7480 | **2.0467** | **351.0062** | **184.222** | **R > S ***** | **2.88** |
| **324;184** | 1.7384 | **1.2634** | **323.9889** | **183.797** | **R > S ***** | **106.8** |
| **280;184** | 1.7056 | **1.2289** | **280.0842** | **183.768** | **R > S ***** | **5.9** |
| **432;184** | 1.6488 | **1.8309** | **431.9707** | **183.776** | **R > S ***** | **3.44** |
| **86;184** | 1.6112 | **1.0146** | **86.05939** | **184.278** | **R > S ***** | **1.87** |

^1^ When undetermined, VIP are identified through m/z;rt values. VIP values are classified in decreasing order.

^2^ mass to charge ratio

^3^ retention time

^4^ The significance was assessed through a Wilcoxon test at *** *P* < 0.001, ** 0.001 < P < 0.01
